# Climate associated natural selection in the human mitochondrial genome

**DOI:** 10.1101/2025.05.23.655374

**Authors:** Finley Grover-Thomas, Lucy van Dorp, Francois Balloux, Aida M. Andrés, M. Florencia Camus

## Abstract

Mitochondria are essential for cellular energy production and biosynthesis, thermogenesis, and cell signalling, and thus help coordinate physiological responses to changing environments. Humans (*Homo sapiens*) have adapted to cope with a wide range of climatic conditions, however the role of the mitochondrial genome (mtDNA) in mediating this process remains poorly understood. Here we curated a dataset of 23,790 publicly available full human mitochondrial genomes, an approximate 40-fold increase on earlier studies, paired with modern climate and reconstructed paleoclimate variables. Using a Generalised Linear Model approach, we identify 18 independent candidate variants significantly associated with climatic conditions, suggesting local adaptation in human mitochondrial genomes. Candidate variants are distributed across multiple loci in regulatory, tRNA, rRNA and protein-coding regions - including prominently in *ND2* and *ND4* complex I subunits. Specific variants are predicted to impact mtDNA transcription, ribosome or protein structure, and multiple have been associated with disease pathologies. We further show that candidate variant genotype distributions are each best modelled by different paleo-bioclimatic variables, consistent with environmental stressors linked to our measured variables exerting subtly distinct selective effects. These stressors may reflect dietary changes or different thermogenic demands at lower temperatures. Our results provide genetic evidence to support the accumulating body of work from functional studies that mitochondria can modulate adaptation to diverse environments. This work underscores the importance of mtDNA in evolutionary biology and its relevance for understanding both disease and physiological variation in global populations.

## Introduction

Mitochondria are integral to eukaryotic life, with efficient mitochondrial function critical for cellular processes and many aspects of life-history trait evolution (Spinelli and Haigis 2018). Responsible for ATP synthesis via oxidative phosphorylation (OXPHOS), these organelles of endosymbiotic origin (Gray 2012; Roger, Muñoz-Gómez, and Kamikawa 2017; Zimorski et al. 2014) sit at the centre of metabolic pathways and are directly responsible for the conversion of food-derived substrates into usable cellular energy (Fernie, Carrari, and Sweetlove 2004; Hatefi 1985; Mitchell 1961; Osellame, Blacker, and Duchen 2012). These complex metabolic processes involve the coordinated action of proteins encoded by both the nuclear and mitochondrial genomes (Chinnery and Hudson 2013), the latter of which contains a small contingent of 13 protein coding regions, 2 rRNAs, 22 tRNAs, and a small regulatory region (for a more detailed description of mitochondrial DNA, see supplementary information). Additionally, OXPHOS impacts thermogenesis (Chouchani, Kazak, and Spiegelman 2019; Chung and Schulte 2020) and a multitude of cell signalling pathways (Papa et al. 2012; Spinelli and Haigis 2018; Zhivotovsky et al. 1998), which are critical components of physiological responses to environmental change.

Despite their functional relevance, the use of mitochondrial DNA (mtDNA) in population genetic studies has largely been limited to studies of demographic history, with common mtDNA variants classically used as informative markers of neutral evolutionary processes. This approach has been predominately driven by the practical advantages of sequencing a genome of small size and high copy number, which has negligible signatures of recombination (in most animals) (Wilson et al. 1985). The vital importance of mtDNA and mitochondrial function for individual fitness suggests, however, that beneficial variants could have significant fitness effects, and thus may increase in frequency within a population due to natural selection. Over the past two decades, a growing body of experimental and statistical genomic work has provided evidence for selection acting on the mtDNA genome of both plants and animals (Awadi et al. 2021; Consuegra et al. 2015; Foote et al. 2011; Garvin, Bielawski, and Gharrett 2011; Havird et al. 2017; Kan, Liao, and Wu 2022; Lamb et al. 2018; Noll et al. 2022; Yu et al. 2011; X. Zhang et al. 2024). Furthermore, clines in mtDNA haplotype frequencies have been detected within several species, potentially indicating signatures of adaptation (Camus et al. 2017; Silva et al. 2014). These results suggest a more active role of the mtDNA in evolutionary adaptation than classically assumed.

In parallel, laboratory studies have examined the functional relationship between environmental factors and mitochondrial function. These studies have demonstrated that both life-history traits (Aw et al. 2018; Hoekstra, Siddiq, and Montooth 2013; Lajbner et al. 2018) and the inheritance patterns of mtDNA can be influenced by thermal and dietary landscapes (Doi, Suzuki, and Etsuko T. Matsuura 1999; E. T. Matsuura, Tanaka, and Yamamoto 1997). Such studies have typically relied on experimental designs that compared mitochondrial haplotypes from highly diverged populations or distinct species. These experimental designs maximise the opportunity to detect mitochondrial genetic effects due to increased mitochondrial divergence but are less informative of the consequences of intra-species mtDNA diversity. Nevertheless, they suggest that variation in the mtDNA is responsive to environmental selection, at least in laboratory settings.

Anatomically modern humans (AMH, *Homo sapiens*) are of particular interest, given that we have successfully colonised a diverse range of environments: from boreal forests to arid desserts and tropical rainforests. Exposure to many of these environments has occurred in the relatively recent past (Betti et al. 2020), following the out-of-Africa expansion 50,000-100,000 years ago (Beyer, Krapp, Eriksson, et al. 2021; Prugnolle, Manica, and Balloux 2005). These expansions predominately occurred at times of large climate fluctuations (35), with many demographic movements across all habitable continents continuing after the last glacial maximum approximately 20,000 years ago. Adaptation to these divergent environments will have required metabolic flexibility to modulate cellular energetics and metabolic processes, possibly impacting core physiological functions including cold-induced thermogenesis, heat dissipation, and metabolic efficiency across different diets. Mitochondrial DNA, encoding the core components of the mitochondrial machinery (Chinnery and Hudson 2013; Chinnery, Mowbray, et al. 2007), may thus encode mutations that mediate bioenergetic trade-offs between en-ergy production, biosynthesis, and thermal regulation, and are therefore strong candidates for adaptation to different climates in human populations.

The potential role of mitochondria in mediating human adaptation to these different environments was debated in the early 2000s. The first studies examining this hypothesis using molecular biology and sequencing techniques revealed patterns of amino acid changes which could be associated with broad climatic zones (Mishmar et al. 2003; Ruiz-Pesini et al. 2004). These studies were followed by the discovery of a correlation between the genetic divergence of human mitochondrial haplotypes and temperature differences across human populations (Balloux et al. 2009). However, studies using different datasets did not all reach the same conclusions (Kivisild et al. 2006; Sun, Kong, and Y.-P. Zhang 2007). In the last 15 years, the number of mtDNAs sequenced has hugely increased, and over 60,000 human mitochondrial genomes are now publicly available on NCBI at the time of writing –a forty-fold increase over previous studies. The increased genomic datasets, together with a parallel expansion in datasets mapping climatic, biodiversity, and disease outbreak variables geographically, advances in population genetics methods, and an improved understanding of mitochondrial physiology and its genetic variants, makes it timely to revisit the global distribution of human mitochondrial genotypes with respect to the environment. Here, we capitalise on these advancements to test the potential influence of local genetic adaptation in the evolution of human mitochondria, leveraging an extensive dataset of over 23,000 published human mitochondrial genomes spanning 80 countries distributed globally. Co-analysing mtDNA sequences alongside proxies for climate we test whether mtDNA variants have likely contributed to local adaptation in humans and, if so, where they are located, and which climatic variables are the strongest proxies for selective drivers.

## Methods

Unless otherwise specified, computational analysis was performed using R Statistical Software version 4.3.0 (R Core Team 2023), and plotted using the *ggplot2* library (Wickham 2016).

### Data curation

All publicly available GenBank files for sequences matching the search term “(00000015400[SLEN]: 00000016600[SLEN]) AND Homo[Organism] AND mitochondrion [FILT] AND (15400[SLEN] :17000[SLEN])” were downloaded from NCBI using the *rentrez* (Winter 2017) R package. This search term is part of the query provided by the MITOMAP database compendium (Lott et al. 2013) and ensured that only full-length mitochondrial sequences were downloaded. Accession numbers, organism name, country and notes categories, cell line names, and full sequence information was extracted using a custom Python script. Sequences deriving from cell lines and ancient hominids were removed, and country names were paired with country codes in accordance with the ISO 3166 International standards. All analysis was then performed at the country scale. This reduced the geographical resolution of the data in some instances but ensured consistent reporting across data points (countries). The resulting dataset considered 23,850 sequences, an approximate 40-fold increase compared to earlier studies.

For each country mean latitudes and areas were calculated using the *geodata* (Hijmans et al. 2023) and *terra* (Hijmans 2023) packages in R. Latitude was then used as a proxy (Supplementary Figure 1) for temperature-related bioclimatic variables in WorldClim 2.1 (Fick and Hijmans 2017). Annual precipitation (BIO12) was extracted from the WorldClim 2.1 dataset at a spatial resolution of 2.5 minutes (approximately 21km^2^ at the equator) and averaged per country. Whilst latitude was used here in the first stage of analysis, as in earlier studies (Balloux et al. 2009; Key et al. 2018), reconstructed ‘biologically relevant’ paleoclimatic variables may serve as more specific proxies for ancestral environmental conditions. Paleoclimate variables reconstructed by Beyer *et al* (Beyer, Krapp, and Manica 2020) were downloaded using the *pastclim* (Leonardi et al. 2023) R package and a set of 10 bioclimatic variables (Table 1) were selected for consideration. “Mean temperature of the driest quarter” and “Precipitation of the driest quarter” have both previously been associated with ancient human migration routes (Beyer, Krapp, Eriksson, et al. 2021), and are used here as proxies for freshwater availability and diet, as is ’leaf area index’. The ‘temperature of the wettest quarter’, ‘mean temperature of the coldest quarter’, and ‘mean temperature of the warmest quarter’ may in turn reflect potentially challenging physiological environmental conditions. Paleoclimate variables were averaged over the last 20,000 years and per country. This averaging approach yielded mean values for both modern and paleoclimate variables that correlated strongly with the extracted values from country centroid locations and capital city coordinates (Supplementary file 2).

**Table 1:** Bioclimatic variables. Reconstructed paleo-bioclimatic variables (and corresponding WorldClim identifier) obtained and tested for associations with mitochondrial DNA SNP variants.

| Abbreviation | Name | Units |
| --- | --- | --- |
| bio1 | Mean annual temperature | °C |
| bio4 | Temperature seasonality | °C |
| bio8 | Mean temperature of the wettest quarter | °C |
| bio9 | Mean temperature of the driest quarter | °C |
| bio10 | Mean temperature of the warmest quarter | °C |
| bio11 | Mean temperature of the coldest quarter | °C |
| bio12 | Annual precipitation | mm / year |
| bio17 | Precipitation of the driest quarter | mm / quarter |
| bio19 | Precipitation of coldest quarter | mm / quarter |
| lai | Leaf Area Index | gC per $m^2$ |

We additionally approximated patterns of plant biodiversity and pathogen load by region, using data from the World Checklist of Vascular Plants (WCVP) (Brown et al. 2023) and Torres Munguía et al. 2022 respectively. Botanical countries were grouped by the corresponding ISO 3166 country names, then the mean number of endemic species and outbreaks per country were calculated and normalised by geographical area.

All sequences were sequentially aligned to the Cambridge Reference full human mitochondrial genome (rCRS, NC012920.1) using MAFFT v7.490 software (Katoh and Standley 2013). Visual inspection in UGENE (*Unipro UGENE: a unified bioinformatics toolkit | Bioinformatics | Oxford Academic* 2025) identified spurious or misaligned sequences. To automate the curation of the alignment, a custom R script was used to remove sequences that inserted a gap into more than 99.97% of sequences, as these were considered likely sequencing or alignment errors. Remaining sequences were realigned to the reference genome and this process was repeated until no insertions were present. The final alignment contained 23,791 sequences and was converted to a VCF format using the *snp-sites Bioconda* (Page et al. 2016) package. A minor allele frequency cut-off of 0.005% was applied using *VCFtools* (Danecek et al. 2011) and site frequency spectra distributions were visually inspected. The resulting VCF file contained genotype information for 422 variable sites and was imported into R using the *ape* package (Paradis and Schliep 2019).

A maximum likelihood phylogenetic tree was reconstructed for all 23,790 sequences (reference individual duplicate removed) using *FastTree2* software, version 2.1.11 (Price, Dehal, and Arkin 2010). Data visualisation in *Taxonium* (Sanderson 2022) and R showed that these sequences did not cluster by country and there were many recurrent mutations across distinct branches. Principal component analysis (PCA) was run using the *adegenet* R package (Jombart 2008) on the same complement of sequences. Countries with fewer than 20 total individuals were removed from downstream analyses. For the four countries with more than 1000 individuals (N_ESP_= 2478; N_RUS_ = 2358; N_GBR_ = 1845; N_USA_ = 1053) 1000 individuals were randomly selected. The final collection of sequences thus contained 19,570 full mitochondrial genomes from 80 countries.

### Latitude and ‘Modern’ Annual Precipitation GLMMs

Prior work by Key and colleagues demonstrated that Generalised Linear Mixed Models (GLMMs) can be used to help infer temperature-related positive selection in humans (Key et al. 2018). Whilst that research studied a nuclear gene (*TRMP8*), we propose that this approach is also suitable for application to human mtDNA sequences.

Genotypes (reference/alternative) were modelled separately for each SNP as a binary response variable with a binomial error distribution using the *lme4* R package (Bates et al. 2015). Principal components 1 to 4, for each individual, were included as fixed effects in each model to capture patterns of shared ancestry between individuals. The associated country was used as a random intercept to account for the non-independence of shared ancestry and environmental effects within countries. Models with the principal components and the random intercepts only were considered as the null model. Given the complexity of the models, the Bound Optimisation By Quadratic Approximation method was used with the maximum number of iterations increased to 100,000 to improve the likelihood of convergence. SNPs where the null model did not converge after this number of iterations were removed alongside models with a dispersion ratio below 0.8. These were typically sites with very low allele frequencies. 203 SNPs remained after this filtering stage for further analysis.

We next investigated the association of the environmental variables to each mtDNA SNP genotype. To do so we included latitude and annual precipitation (WorldClim, BIOL12) as joint test predictors with fixed effects in an alternative model (“LatPrecip GLMM”) and compared to the null using a Likelihood Ratio Test (LRT). A conservative 1% alpha significance threshold was corrected using the Bonferroni calculation to account for multiple testing across sites and minimise the risk of false positive results. Principal components were sequentially dropped from the alternative model and LRTs were used to assess the associated changes in model accuracy. SNPs where the distribution of genotypes were significantly better explained by the alternative than the null model were designated as “Latitude and Precipitation candidate variants” (hereafter “LatPrecip candidate variants”). Where the Pearson’s R^2^ correlation coefficient between multiple LatPrecip candidate variant genotypes was greater than 0.2, the associated variants were clustered together. The most significant variant was then selected as the representative “LatPrecip candidate variant” for the cluster. Correlation coefficients and clustering were calculated using the *Hmisc* (Jr 2024) and *stats* (R Core Team 2023) R packages respectively.

### Paleo-bioclimatic and Disease Predictor GLMMs

In the paleo-bioclimatic analyses, Latitude and (modern) annual precipitation were replaced as predictors in the alternative model by either one of the 10 paleo-bioclimatic variables, the number of endemic plant species, or the number of disease outbreaks. New alternative models (“PaleoBioGLMMs”) were fitted for each of the new predictors for all 203 SNPs which had passed the earlier null model convergence and dispersion filtering steps. All PaleoBio-GLMMs were compared to the existing null model using the LRT as before, however a different Bonferroni-corrected threshold was used to account for the additional 12 new GLMM predictors tested. For SNPs where at least one of the Paleo-GLMMs significantly outperformed the null model, the SNP was labelled as a “PaleoBio candidate variant” and the predictor used was assigned as the “significant bioclimatic predictor”. For SNPs where multiple PaleoBio-GLMMs (each with a different predictors) surpassed the significance threshold, the strongest (i.e.: most significant) predictor was designated the “significant bioclimatic predictor” for that SNP. We acknowledge this approach does not account for the magnitude of the difference between the best and second-best predictors for a given site. As with the LatPrecip GLMM analysis detailed above, variants with an R^2^ Pearson’s correlation coefficient between genotypes above 0.2 were clustered and the strongest signal was selected as the representative candidate for the cluster.

### Candidate Loci Positions and Functional Effects

We examined the possible functional effects of candidate loci detected within protein coding regions using ENSEMBL’s Variant Effect Predictor (McLaren et al. 2016). This toolset uses existing collections of genomic annotations to determine the likely impact of a variant on structure and phenotype, and classifies a variant as benign, deleterious, or unknown using the SIFT and PolyPhen scores. For candidate loci within tRNA-encoding sequences, the *tRNAScan* online software tool (Chan and Lowe 2019) was used to predict secondary structures.

### TreeWAS phylogenetic analysis

Tree-based phylogenetic approaches have also been used to assess the potential association between genotypes and environmental variables (analysed as “phenotypes”) while accounting for shared ancestry. *TreeWAS*, originally designed for use on bacterial sequences with high mutation rates, is one such approach that identifies systematic differences in genotypes across a phylogenetic tree that correspond to “phenotypic” differences while accounting for clonal lineage structures (Collins and Didelot 2018). TreeWAS’ ‘subsequent test’ reconstructs ancestral phenotypes and genotypes for each node of a phylogenetic tree, and calculates in what proportion of tree branches the genotype and phenotype are expected to be in the same state (Collins and Didelot 2018). To test correlations between genotypes and environmental vari-ables in an independent analysis, we ran a TreeWAS test using latitude as the phenotype using the *TreeWAS* R package. We used a pruned version of the FastTree2 phylogenetic tree which only contained the 19,570 sequences used in the LatPrecip and PaleoBio GLMM analyses. A limitation of this approach, and a difference with the GLMM analysis, is that only one variable (latitude) can be tested at a given time. To compare the results of the GLMM analyses and TreeWAS, we categorised SNPs into those that correlated with LatPrecip candidate variants in the GLMM (Pearson’s R^2^ correlation coefficient *>* 0.5) and those that did not (no GLMM correlation observed); the distributions of TreeWAS p-values were then compared between the two groups using a Wilcox rank sum test.

## Results & Discussion

We curated a dataset of 19,570 human mitochondrial DNA sequences from 80 countries. Assessing variation over 203 mtDNA SNPs, we tested for associations between mtDNA genotypes and a range of climatic variables. Such associations would be consistent with mtDNA mediating adaption to the diverse range of environments that humans inhabit.

### Identification of Candidate Variants

If local adaptation to climate has shaped the evolution of the mtDNA in humans, we would expect that the frequencies of mtDNA variants would correlate with climatic variables more than expected under neutrality. To test this hypothesis, we first assessed if climate proxies are significant predictors of mtDNA SNP genotype distributions. A Generalised Linear Mixed Model (GLMM) approach was used to test for associations between two climatic variables (latitude and annual precipitation) and the genotypes of each mtDNA SNP while accounting for shared ancestry between individuals. In the absence of associated nuclear genomes, and given the different demographic history of the nuclear and mitochondrial genomes, the mitochondrial genomes are used to both account for demography and to test for selection.

Our results show that at least one principal component (derived from all SNPs and used to account for shared ancestry) was a significant predictor (*α* = 0.01, *p* < 8.25 *×* 10^−6^) for 96.5% of SNPs tested, including every candidate variant. This indicates that our estimates of shared ancestry do adequately capture and explain the genetic variance expected under neutrality for the vast majority of SNPs (see Supplementary detail for more information). For the majority (85%) of SNPs tested, the alternative model containing latitude and annual precipitation as additional predictors failed to significantly outperform the null model with shared ancestry alone (*α* = 0.01*, p >* 4.9 *×* 10^5^, Figure 1), consistent with neutral demographic processes having strongly shaped the distributions of mtDNA genotypes. Nevertheless, in 18 SNPs, the null model with shared ancestry alone was significantly outperformed by the alternate model containing latitude and annual precipitation as additional predictors (*α* = 0.01*, p <* 4.9 *×* 10^−5^) (Figure 1). We refer to these SNPs as “Latitude Precipitation (“LatPrecip”) candidate variants”.

**Figure 1:**
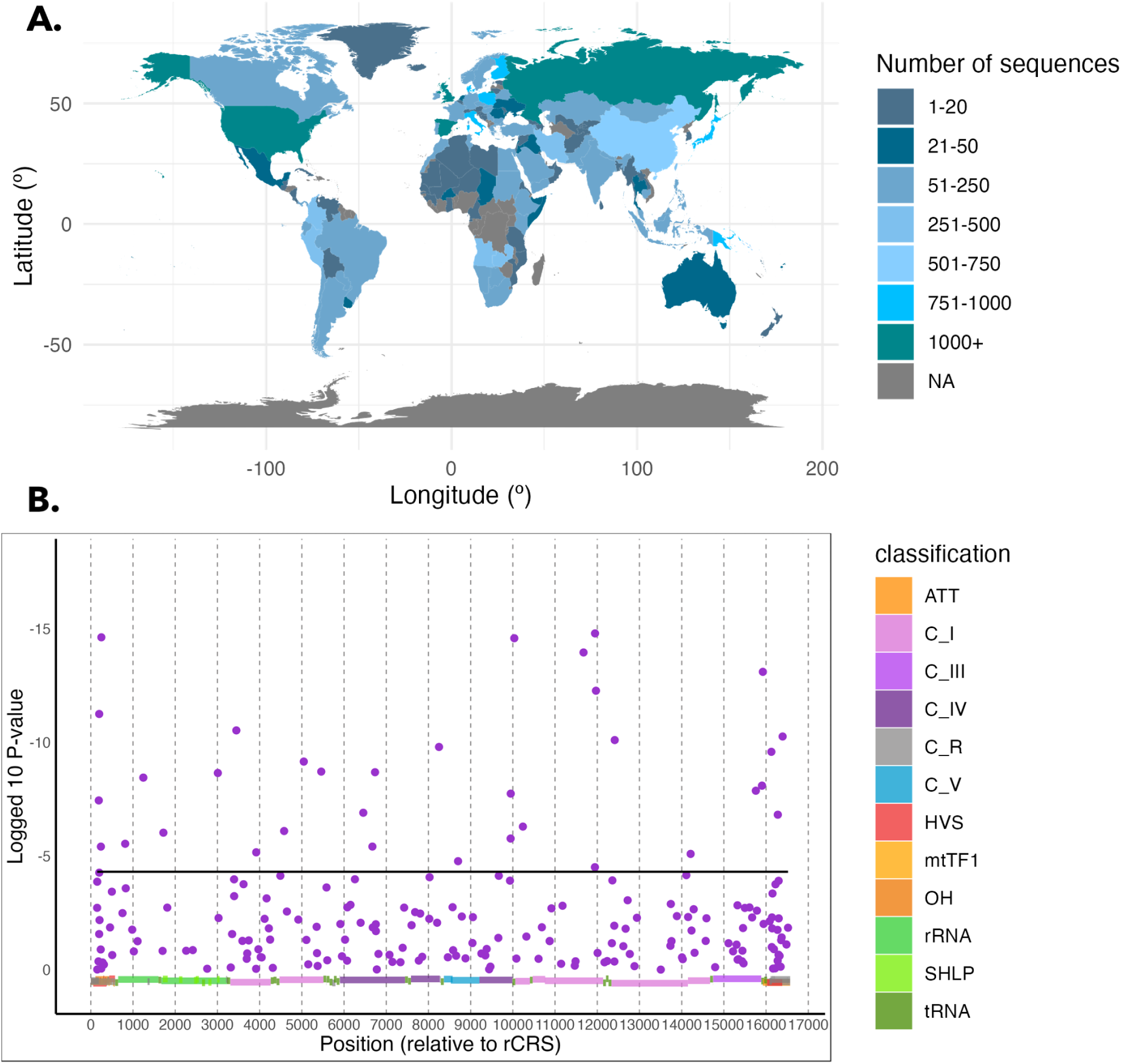
(**A. Global distribution of sequences.** Colours indicate the number of sequences per country in the curated dataset. NA indicates no full mtDNA sequences retrieved. Sequences from countries with fewer than 20 total sequences (grey-blue) were not included in GLMM anlaysis. **B. Climate associations of mtDNA variants** Log_10_ p-values from likelihood ratio tests comparing GLMMs with and without climate variables (latitude and annual precipitation). Purple dots indicate a score for each of the 209 SNPs tested. The black solid horizontal line denotes the Bonferroni-corrected 1% significance threshold. Coloured blocks show loci classification (red & orange: control region; light green: rRNA subunits; dark green: tRNAs; light pink: complex I subunits; light purple: cytB; dark purple: complex IV subunits; blue: complex V subunits).

In the absence of recombination, a reasonable concern is whether these represent 18 separate signals. Our results provide evidence that they do. First, candidate variants are not highly correlated with each other (Supplementary Figure 2). This is as, when multiple significant SNPs are in linkage (Pearson’s R^2^ correlation coefficient *≥* 0.2, see methods for more details), only the single most significant SNP is designated as a candidate variant. Second, all LatPrecip candidate variants re-occur in multiple, separate branches of the phylogenetic tree (Supplementary Figure 3). This reduces the likelihood that the variants are linked by shared ancestry and suggests instead independent selective pressures in multiple lineages. The presence of these 18 (“LatPrecip”) candidate variants represent a 28-fold increase in the number of significant SNPs relative to the number expected by chance alone (see Supplementary Information). Furthermore, removing sequences from the Americas, Australia, and New Zealand did not affect the results (Supplementary Information, Supplementary Figure 4), suggesting that the observed signals are not artefacts from recent migration and admixture. For 200 0f 203 SNPs, adding the first environmental principal component (derived from a PCA of 11 bioclimatic variables) to the model failed to significantly improve model fit relative to the alternative model with latitude and annual precipitation only (Supplementary Figure 5), indicating that latitude and annual precipitation themselves capture relevant variation in climatic conditions.

These results are also supported by an alternative phylogenetic approach, TreeWAS (Collins and Didelot 2018). SNPs in linkage (R^2^ *≥* 0.5) with LatPrecip candidate variants had TreeWAS p-values that skewed towards zero (Wilcox rank-sum test, W = 1099, p < 0.001, Supplementary Figure 6), suggesting that LatPrecip candidates also have lower p-values in TreeWAS. No SNPs passed the Bonferroni-corrected threshold in the TreeWAS analysis, this is likely because this method has been designed and optimised for bacterial sequences with a higher mutation rate and well-resolved trees devoid of polytomies. Additionally, only temperature could be accounted for in TreeWAS, as only a single phenotypic trait can be supplied in the existing approach.

There was a strong bias among LatPrecip candidate variants for SNPs with higher minor allele frequencies (MAFs) at higher latitudes. This is consistent with the patterns expected from local adaptation, however this may have been exaggerated by an under-representation of African and South American sequences (Supplementary Information), reducing the likelihood that recurrent mutations would be identified in these populations. Increasing the number of publicly available genomic sequences from the global south is important for equity and parity. Genomes from the global south contain unique genetic variants (Fortes-Lima and Schlebusch 2021; Merriwether et al. 1991), and inhabit a larger variety of environmental conditions, including at the within-country scale. Greater sampling in Africa may also reveal additional sites and variables of interest related to more ancestral climate adaptations, which may be masked by more recent adaption to the physiological demands of novel environments in Europe and Asia.

### Distribution Candidate Loci

The 18 LatPrecip candidate variants are dispersed across 21 loci, with four variants (m.189, m.250, m.3010, and m.16126) overlapping multiple functional regions, as visualised in Figure 2. Three complex I subunits (*ND3, ND4L, and ND5*), complex III subunit II (*COII*), and *ATP synthase subunit 8* (*ATP8*) are devoid of any LatPrecip candidate variants. Eleven variants are found in protein-coding regions, while seven are not. Two (m.10034 and m.15904) are within mtDNA transferRNAs (tRNA-glycine and tRNA-threonine respectively), two are within the ribosomal RNA loci (m.813, 12S; and m.3010,16S), and three are within the control region (m.189, m.250, and m.16126). The four most significant candidates in protein-coding regions (in order of decreasing significance: m.11947, synonymous; m.11969, missense; m.3447, synonymous; m.5460, missense) are all located in subunits of complex I, and the two most significant are located within *NADH dehydrogenase subunit 4* (*ND4*). These are m.11969, previously identified as a conserved root variant for Siberian haplogroup C with a possible role in cold adaptation (Ruiz-Pesini et al. 2004), and m.11947, the top LatPrecip candidate, which has not previously been associated with environmental adaptation.

**Figure 2:**
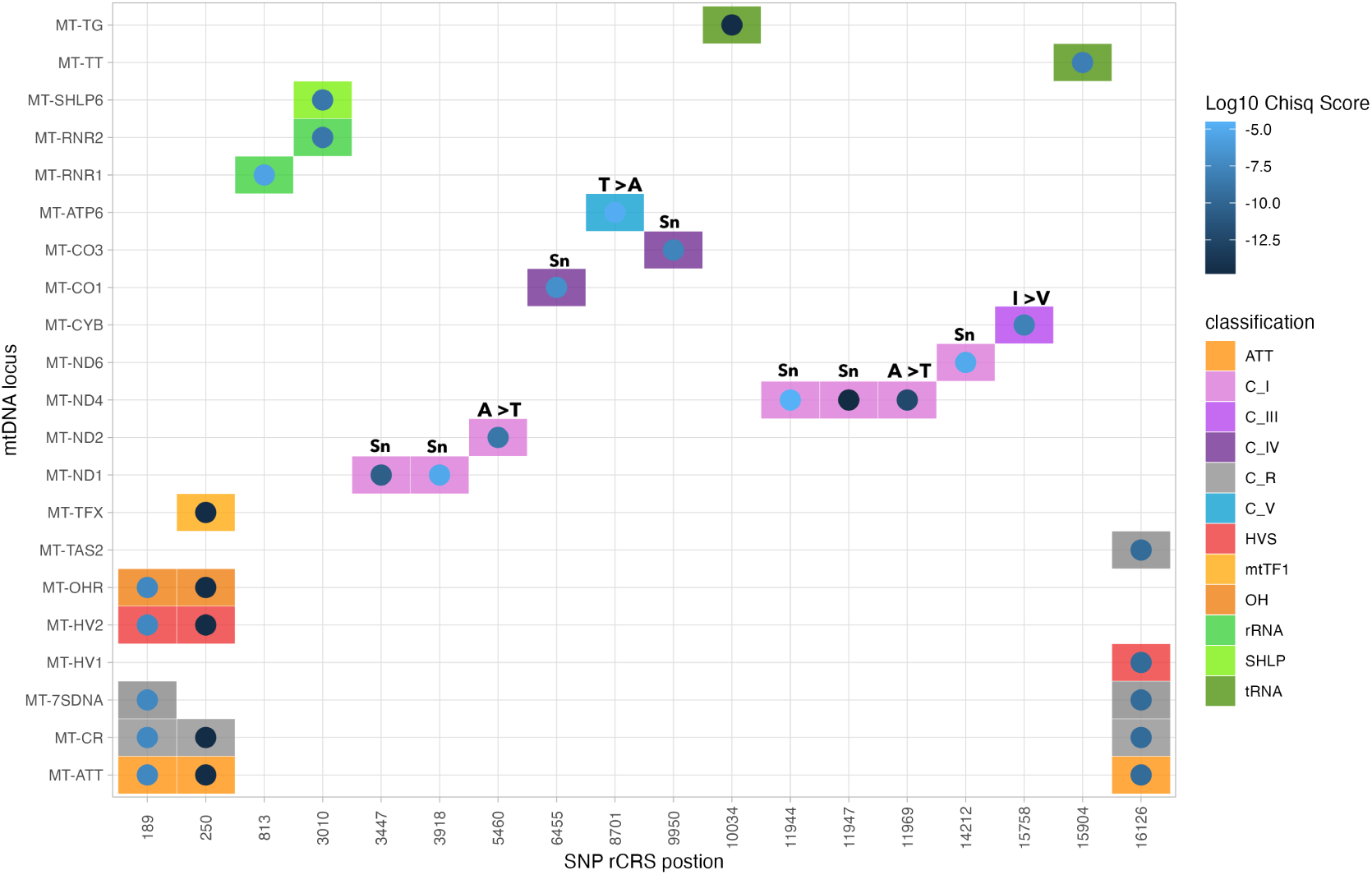
Positions of candidate loci across the mtDNA Candidate SNPs displayed vs mtDNA loci. Log_10_ p-values from LRT (dot colour) and MITOMAP locus classification (tile colour) are additionally represented. (ATT: membrane attachment site, indicates a role in mtDNA replication; C I: complex I; C III: complex III; C IV: complex IV; C R: control region; C V: ATP synthase; HVS: hypervariable sequence; mtTF1: mtDNA transcription factor 1 binding site; OH: origin of H-strand replication; rRNA: ribosomal RNA subunits; SHLP: short humanin-like proteins; tRNA: transfer RNA). Annotations above coloured tiles represent whether protein-coding variants are synonymous (Sn) or missense (single letter amino acid code changes) changes.

The strongest signatures in protein coding genes were thus observed for SNPs in the *ND4*, *ND1* and *ND2* subunits of complex I. This is the largest OXPHOS complex, the earliest entry-point for electrons into the electron transport chain (ETC), and the largest producer of reactive oxygen species (ROS) (Chinnery and Hudson 2013; Wolstenholme 1992). Localisation of candidate variants to complex I subunits is consistent with its increased physiological importance relative to the other ETC complexes (Esposti 2017; Scialò et al. 2020). Additionally, our results show striking concordance with studies in salmon, hares, penguins and monkeys that show signatures of local adaptation to the environment in mitochondrially-encoded ND1, ND2 and ND4 protein subunits (Awadi et al. 2021; Consuegra et al. 2015; Garvin, Bielawski, and Gharrett 2011; Yu et al. 2011). These signatures include elevated *d* N/*d* S ratios, elevated numbers of non-synonymous changes within codons across multiple taxa, and enrichment of radical changes in terms of amino-acid physiochemical properties. The observed phylogenetic concordance across a range of species suggests that mtDNA-related mechanisms of adaptation may be conserved across species. These mtDNA genes encode complex I subunits (principally the adjacent *ND2* and *ND4*) and play key roles in the passage of protons into the intermembrane space (Baradaran et al. 2013; Kampjut and Sazanov 2022). A possible mechanism of adaptation is that mutations decreasing the efficiency of this process lead to increased intra-cellular heat production via increased proton leakage, potentially providing an advantage in low temperature environments, but a disadvantage at warmer temperatures.

### Functional Analyses of Candidate Variants

Besides the two missense mutations in *ND2* and *ND4* mentioned above, two more LatPrecip candidate variants are missense mutations. One within *Cytochrome b* (m.15758), and one within *ATP synthase subunit 6* (m.8701), for which the difference in allele frequencies between populations have previously been shown to associate with minimum temperature beyond levels expected from isolation by distance alone (Balloux et al. 2009). Seven candidate variants are synonymous mutations, although experimental work has shown that synonymous mutations in mtDNA can alter mitochondrial physiology and individual metabolic fitness (Bettinazzi et al. 2024; Camus et al. 2017), potentially via a mechanism of overlapping open reading frames, as discussed in more detail below. It thus remains possible that all identified candidate variants in protein-coding regions have functional consequences, even if we cannot currently predict the impact.

Outside of protein-coding genes, two LatPrecip candidate variants are predicted to alter the function of two mitochondrial tRNAs, two are predicted to impact the mitochondrial ribosome, and three may impact mtDNA regulation. Two strong candidate variants (m.15904 & m.10034) are predicted to alter tRNA-threonine’s D-loop and tRNA-glycine’s variable loop respectively (Supplementary Information). *In silico* modelling predicts the m.3010 candidate variant to lower the molar free energy of *16S* rRNA and disrupt the secondary and tertiary structure of the mito-ribosome (Rovcanin et al. 2020); with previous studies linking this variant to cyclic vomiting syndrome, glioblastomas, and migraines (Boles et al. 2015; Rovcanin et al. 2020). We note that the *TRMP8* nuclear variant inferring cold resistance identified by Key et al (2018) was also linked to migraine prevalence, potentially highlighting a fundamental trade-off between efficient thermogenesis and neurological function.

Internal regulation of mtDNA replication and gene expression appears to be a critical, yet often overlooked, factor in how mitochondria mediate adaptation to environmental conditions (Coskun, Ruiz-Pesini, and Wallace 2003). This is supported in our data by the presence of strong candidate variants in key regulatory regions — including the termination-associated sequence TAS2 (m.16126), the origin of heavy-strand replication (m.189) and the Transcription Factor A Mitochondrial (MtTF1) binding site (m.250), as well as the candidate variant m.3010 located within the rRNA. Previous work has identified elevated basal metabolic rates in circumpolar populations (Leonard et al. 2002) and increased mtDNA content in Chinese populations occupying colder northern environments (Cheng et al. 2013); our results suggest directly for the first time that this may be regulated not only by the large nuclear genome, but locally within the mtDNA itself. The comparatively strong performance of paleo-annual precipitation as a model predictor, relative to temperature-related variables for m.16126, as discussed in more detail below, suggests that regulation of mtDNA transcription and translation is likely not only a valuable response to thermogenesis-related cellular demands, but also to other changes in the environment, potentially diet. As with temperature, diet composition alters mtDNA copy number in Angus cattle and humans (Bai et al. 2020; Pollicino et al. 2023), and mutations in the termination associated sequence (TAS2), responsible for regulating mtDNA replication, have been associated with type II diabetes (Jiang et al. 2017), although further work is required to establish the molecular basis of this association.

In addition to mtDNA regulation and expression, we predict that the role of peptides encoded in alternate open reading frames may be important, but additionally overlooked, aspects of local adaptation within mitochondria. The m.3010 variant overlaps not only with *RNR-2* (*ribosomal subunit 16S*) but also *Short Humanin Like Protein 6* (*SHLP6*), one of eight short but functional peptides encoded in alternate reading frames within the mtDNA rRNA regions (Cobb et al. 2016). This provides an alternative explanation for the potential role of this variant in adaptation to colder environments, where the alternative allele occurs at elevated frequencies. SHLP6 is the only Humanin-like peptide within the mtDNA that has a highly conserved start and stop codons and amino acid sequences (Gruschus, Morris, and Tjandra 2023). Sequence conservation has been taken as indicative of a functional impact and expression of the transcribed peptide has been associated with retrograde signalling for thermoregulation and lower body temperatures in heterothermic mammals (Emser et al. 2023). Consequently, it remains possible that the m.3010 mutation facilitates adaptation to cold or variable environments through the SHLP6 signalling pathway, although further work is needed to potentially test this hypothesis. The presence of functionally relevant peptides, including SHLP6, reduces confidence in predictions of whether mtDNA variants are truly synonymous, particularly given the large number of recently discovered alternatively encoded peptides across the entirety of the mammalian mtDNA (Kienzle et al. 2023).

### Paleo-climate, plant diversity, and outbreaks as model predictors

Latitude has long been used as a proxy variable for temperature as it correlates with many temperature-related bioclimatic variables. Latitude is not, however, in itself a selective pressure. An outstanding question is which environmental variable(s) may be driving local adaptation and whether the same one(s) may be driving adaptation in the different SNPs. These are challenging questions to answer given the high correlation between all similar variables (eg.: all temperature-related variables). Nevertheless, if all candidate variants have responded to the same underlying selective force, we might expect, when latitude and annual precipitation are replaced as GLMM predictors by new paleo-bioclimate and disease outbreak variables, the same new variable would be the most significant predictor across all SNPs. We do not observe this in our results (Figure 3) and instead identify seven different bioclimatic variables that are the most significant predictor for at least one SNP. This implies that different candidate variants may be responding to subtly different environmental selective pressures, further supporting our conclusion that the 18 LatPrecip candidate variants are distinct from each other.

**Figure 3:**
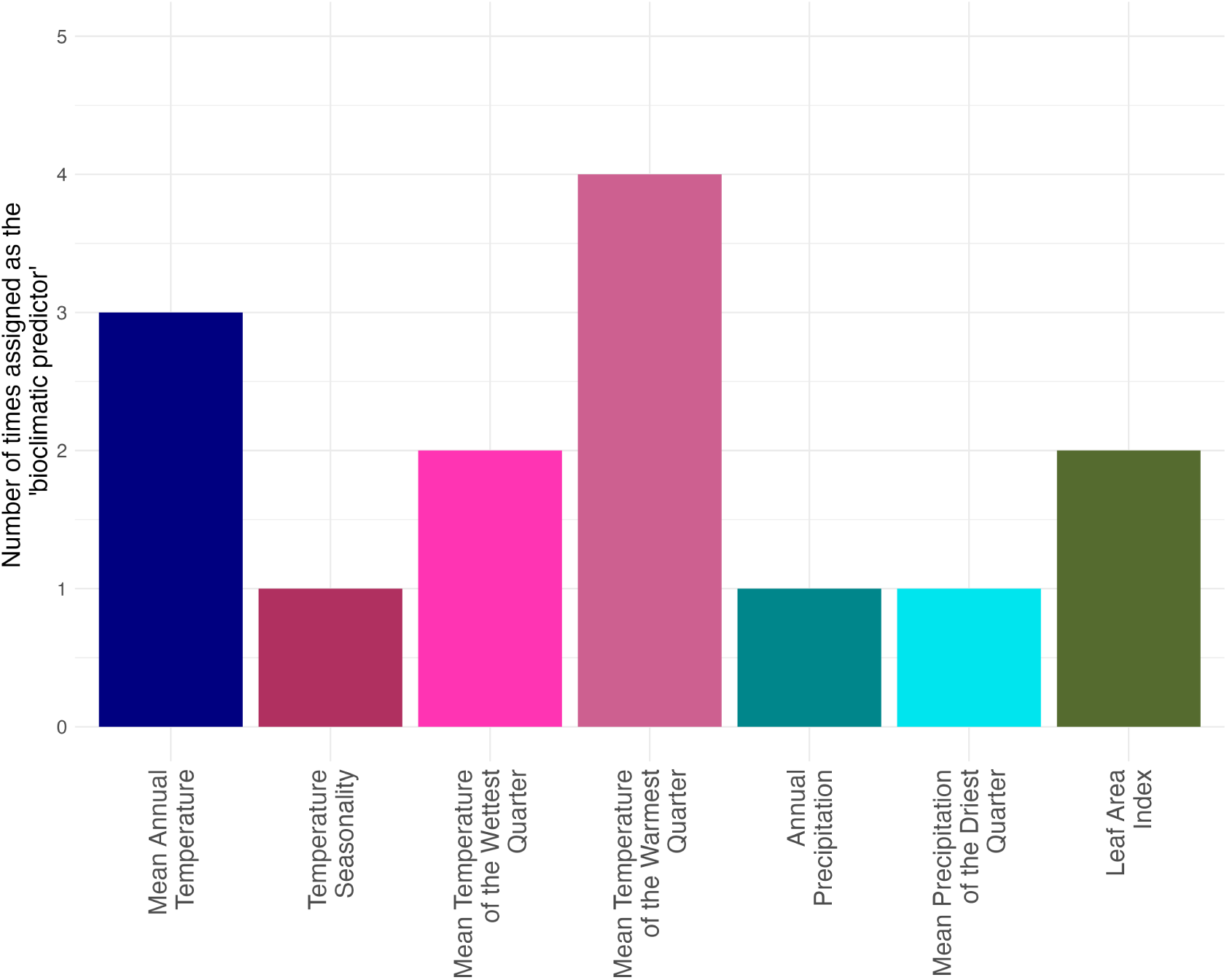
Assignment of ‘bioclimatic predictors’. The number of times a given paleo-bioclimatic variable was both a statistically significant predictor (*α* = 0.01*, p <* 4.11 *×* 10^6^) of a mtDNA SNP genotype in the GLMM analysis and had a lower p-value than any other paleo-bioclimatic predictor tested. Paleo-bioclimatic variables derived from the *pastclim* R package (Leonardi et al. 2023)

We fitted additional GLMMs where fixed effects latitude and WorldClim annual precipitation used above were both replaced in secondary alternative models (PaleoBio-GLMMs, see Methods), by a single paleo-bioclimatic variable, or the number of endemic species, or the number of disease outbreaks. These variables were selected to serve as proxies for different potential selective pressures that relate to mitochondrial function: thermoregulation demands, diets, and disease burden. We identify 14 Paleoclimate candidate variants where the original null model (with shared ancestry alone) was significantly outperformed (*α* = 0.01*, p <* 4.11 *×* 10^6^) by at least one PaleoBio-GLMM. Seven different paleoclimatic variables were assigned as significant bioclimate predictor (i.e.: were included in the PaleoBio-GLMM which most significantly out-performed the null model) (Figure 3). PaleoBio-GLMMs with the number of endemic species and the number of outbreaks did not significantly outperform the null model for any SNPs tested.

Of the seven significant bioclimatic predictors, four included a temperature component, three included a precipitation component, and one (Leaf Area Index) was not a direct estimate of climate. Although it is difficult to draw direct conclusions from these comparisons, they may indicate that the different candidate variants may be mediating responses to subtly different selection pressures. Temperature of the wettest quarter was assigned as the significant bioclimate predictor for two of the top three candidates (Supplementary Data, Table 2) in the PaleoBio-GLMM analysis: m.3010 (Figure 4) and a SNP (m.16391) in strong (R^2^ > 0.5) linkage with m.250, both discussed above in the context of functional effects. Mean temperature of the warmest quarter was commonly assigned as a predictor (with higher MAFs when this temperature decreased, Supplementary Figure 6), consistent with mitochondria potentially mediating adaptation to colder environments which have altered thermoregulation demands. In contrast, the inclusion of precipitation paleoclimate variables and leaf area index as significant bioclimatic predictors suggests that altering thermogenesis is likely not the only mechanism through which mitochondria mediate local adaptation.

**Figure 4:**
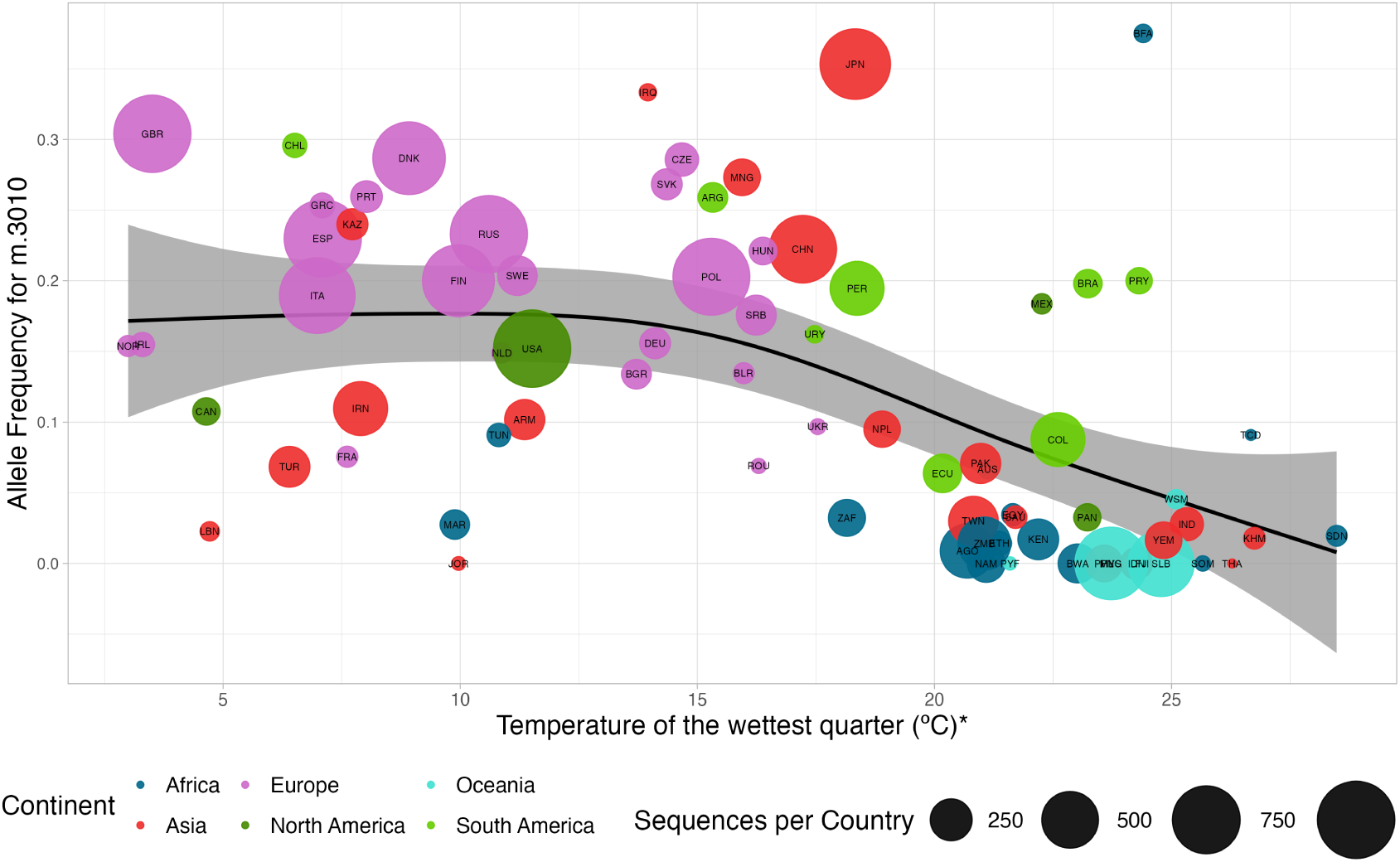
Association between m.3010 derived allele frequency and the temperature of the wettest quarter (°C). Temperature averaged over country area over the last 20,000 years. Circle size represents the number of sequences per country and colour represents the continent (Africa: navy; Asia: red; Europe: pink; North America: dark green; Oceania: turquoise; South America: light green).

**Table 2:**
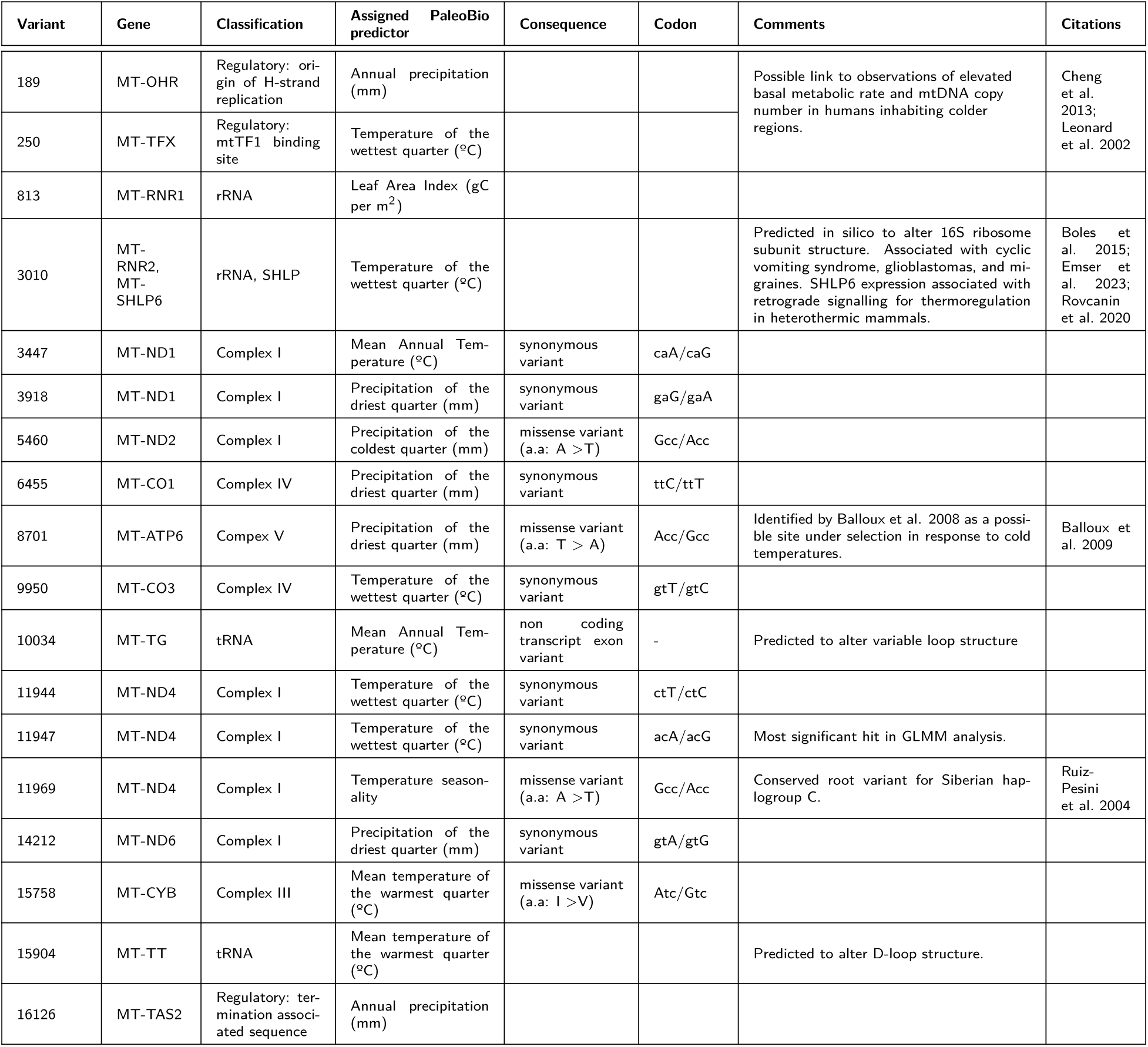
Summary of Candidate Variants. Variant: SNP position relative to the rCRS reference genome; Gene: location of SNP, "*" denotes the SNP overlaps with additional non-coding regions, as described in Supplementary Table 1); Classification: The ETC complex or category of said gene; Assigned Bioclimatic Predictor: the best performing paleo-bioclimatic predictor for the given SNP.

| Variant | Gene | Classification | Assigned PaleoBio predictor | Consequence | Codon | Comments | Citations |
| --- | --- | --- | --- | --- | --- | --- | --- |
| 189 | MT-OHR | Regulatory: origin of H-strand replication | Annual precipitation (mm) |  |  | Possible link to observations of elevated basal metabolic rate and mtDNA copy number in humans inhabiting colder regions. | Cheng et al. 2013; Leonard et al. 2002 |
| 250 | MT-TFX | Regulatory: mtTF1 binding site | Temperature of the wettest quarter (°C) |  |  |  |  |
| 813 | MT-RNR1 | rRNA | Leaf Area Index (gC per m<sup>2</sup>) |  |  |  |  |
| 3010 | MT-RNR2, MT-SHLP6 | rRNA, SHLP | Temperature of the wettest quarter (°C) |  |  | Predicted in silico to alter 16S ribosome subunit structure. Associated with cyclic vomiting syndrome, glioblastomas, and migraines. SHLP6 expression associated with retrograde signalling for thermoregulation in heterothermic mammals. | Boles et al. 2015; Emser et al. 2023; Rovcanin et al. 2020 |
| 3447 | MT-ND1 | Complex I | Mean Annual Temperature (°C) | synonymous variant | caA/caG |  |  |
| 3918 | MT-ND1 | Complex I | Precipitation of the driest quarter (mm) | synonymous variant | gaG/gaA |  |  |
| 5460 | MT-ND2 | Complex I | Precipitation of the coldest quarter (mm) | missense variant (a.a: A > T) | Gcc/Acc |  |  |
| 6455 | MT-CO1 | Complex IV | Precipitation of the driest quarter (mm) | synonymous variant | ttC/ttT |  |  |
| 8701 | MT-ATP6 | Complex V | Precipitation of the driest quarter (mm) | missense variant (a.a: T > A) | Acc/Gcc | Identified by Balloux et al. 2008 as a possible site under selection in response to cold temperatures. | Balloux et al. 2009 |
| 9950 | MT-CO3 | Complex IV | Temperature of the wettest quarter (°C) | synonymous variant | gtT/gtC |  |  |
| 10034 | MT-TG | tRNA | Mean Annual Temperature (°C) | non coding transcript exon variant | - | Predicted to alter variable loop structure |  |
| 11944 | MT-ND4 | Complex I | Temperature of the wettest quarter (°C) | synonymous variant | ctT/ctC |  |  |
| 11947 | MT-ND4 | Complex I | Temperature of the wettest quarter (°C) | synonymous variant | acA/acG | Most significant hit in GLMM analysis. |  |
| 11969 | MT-ND4 | Complex I | Temperature seasonality | missense variant (a.a: A > T) | Gcc/Acc | Conserved root variant for Siberian haplogroup C. | Ruiz-Pesini et al. 2004 |
| 14212 | MT-ND6 | Complex I | Precipitation of the driest quarter (mm) | synonymous variant | gtA/gtG |  |  |
| 15758 | MT-CYB | Complex III | Mean temperature of the warmest quarter (°C) | missense variant (a.a: I > V) | Atc/Gtc |  |  |
| 15904 | MT-TT | tRNA | Mean temperature of the warmest quarter (°C) |  |  | Predicted to alter D-loop structure. |  |
| 16126 | MT-TAS2 | Regulatory: termination associated sequence | Annual precipitation (mm) |  |  |  |  |

An alternate possibility is that mitochondria mediate responses to different diets, in themselves influenced by climatic differences between regions. This possibility is supported by the result that Leaf area index (LAI) was a significant bioclimatic predictor for two variants (m.6671 m.9932) where the minor allele was found at higher frequencies in African and Asian populations with low LAI values (Supplementary Figure 7), indicating barren or semi-barren environments (Parker 2020). Of these two SNPs, m.9932 was not associated with any LatPrecip candidate variant. Given the lower correlation coefficients between latitude and LAI, this is not completely unexpected. No other paleo-bioclimatic predictor, including precipitation measures, were significant for this variant, suggesting that water availability or temperature alone are not driving these signatures. The carrying capacity and resource availability of a region has likely shaped human demography over the last 300,000 years (Beyer, Krapp, Eriksson, et al. 2021), most notably during the out of Africa expansion. It thus remains feasible that LAI may serve as a proxy for changes in diet, and the associations detected here reflect genomic adaptations to this. The localisation of both sites to complex IV may suggest a mechanistic role of complex IV in mediating diet related mtDNA selection, concordant with a link between flavonoids present in red wine and berries and elevated complex IV expression levels (Pollicino et al. 2023), however this line of interpretation remains highly speculative at this stage.

Of the 14 Paleoclimate candidate variants seven are also LatPrecip candidate variants, and an additional four are in strong linkage (R^2^ > 0.5) with LatPrecip candidate variants, demonstrating concordance across tests. The three weakest paleoclimate candidate variants were not, however, correlated with LatPrecip candidate variants, and five of the six weakest LatPrecip candidate variants did not pass the significance threshold after correcting for the additional number of tests in the PaleoBio-GLMMs. While this may have arisen from the conservative thresholds used, it may also reflect that complex ecological interactions are being approximated by a smaller number of variables in the PaleoBio-GLMMs. Latitude and annual precipitation combined will likely capture a greater proportion of the environmental variance than a single paleoclimate variable, thus increasing the likelihood that the alternative model will outperform the null at SNPs mediating adaptive responses to nuanced changes in the environment.

### Mitonuclear coevolution

Efficient mitochondrial function is reliant upon coherent cooperation between the mitochondrial and nuclear genomes, leading to mitonuclear coevolution (Burton and Barreto 2012; Hill 2016; Wolff et al. 2014). Examples of mitonuclear incompatibilities, whereby there is a break-down in this relationship, are being increasingly well characterised (Barreto and Burton 2013; Camus et al. 2017; Lajbner et al. 2018; Moran et al. 2024; Rodríguez et al. 2021; Weaver et al. 2022). Environmental conditions may mask or exacerbate the fitness consequences of small mismatches. Therefore, understanding which climatic variables apply the strongest selective force onto mitochondria may improve understanding of when mitonuclear incompatibilities are most likely to be exposed, valuable in both an ecological (Bettinazzi et al. 2024; Biot-Pelletier et al. 2023; Immonen et al. 2020; Morales et al. 2018) and medical (Almannai, Salah, and El-Hattab 2022; Gershoni et al. 2014) context. The nature of the dataset and approach used in this study, whereby only publicly available mitochondrial genomes were used (which lacked associated nuclear genomes), prevented us from investigating the interaction between mitochondria associated nuclear variants and climate: this should be addressed in future studies.

## Conclusion

TCA cycle metabolism, OXPHOS and ATP synthesis, cell signalling, and thermogenesis can all be optimised by mitochondria, resulting in adaptation to divergent environments across populations (Hill 2016). Climate may exert a selective force via divergent mechanisms, most likely through dietary changes and thermoregulation requirements. Here we show that 18 SNP variants within human mitochondrial genomes are associated with bioclimatic variables at the global scale, highlighting the potential role of mitochondria in human adaptive responses. Our results highlight how treating mtDNA as a neutral marker inhibits our ability to critically evaluate mechanisms of local adaptation at a time where it is particularly important to understand how individuals and species respond to widespread environmental change. Understanding the evolutionary history of adaptive variants may further help identify potential future therapeutic targets in humans and aid ecological understanding of mitochondrial function in a rapidly changing world.

## Acknowledgements

F.G-T thanks Dennis Grover for his support from afar. We thank Max Reuter, Richard Mott, Garrett Hallenthal, and all the members of the Burbano, Murray, Camus and Andrés groups for valuable comments and discussions. We thank all participants who contributed mitochondrial sequences to the original projects, and the authors of said studies for uploading these genomes to the NCBI database. Funding: F.G-T is supported by the UKRI Natural Environment Research Council through the London NERC Doctoral Training Programme scholarship, grant number NE/S007229/1. M.F.C was funded by a UKRI Fellowship (NE/V014307/1) and a Leverhulme Trust grant (RPG-2023-198). L.vD was funded by a UKRI Future Leaders Fellowship (MR/X034828/1) and European Union END-VoC consortium grant (agreement no. 101046314).

## Author Contributions

F.B and M.F.C conceptualised the study; F.G-T, L.vD, F.B, A.M.A and M.F.C designed the analysis. F.G-T collected the data, conducted the analysis, and wrote the original draft; A.M.A and M.F.C supervised the study. All authors contributed substantially to revisions.

## Data Availability Statement

Supportive information is provided as supplementary text and tables. All code and data required to reproduce our analysis is provided at https://github.com/FiG-T/climate_dist_mtDNA.git.

