## Supplementary 1 for "Climate associated natural selection in the human mitochondrial genome"

#### Mitochondrial DNA

The evolution of the mtDNA is strongly influenced by dynamics arising from clonal reproduction, uniparental inheritance solely through the maternal line, as well as negligible rates of recombination but a mutation rate 20-fold (in mammals) higher than the nuclear genome (Brown, George, and Wilson 1979; Lynch, Koskella, and Schaack 2006). The small circular mtDNA genome found in animals typically contains 37 loci: 13 protein coding genes, 22 transfer RNAs (tRNA), and 2 ribosomal RNAs (12S and 16S rRNA). Of the protein coding genes, seven (ND1, ND2, ND3, ND4, ND4L, ND5, ND6) relate to complex I sub-units, one (CytB) relates to a complex III sub-unit, three (COI, COII, COIII) relate to complex IV sub-units, and two relate to complex V sub-units (Björkholm et al. 2015; Chinnery and Hudson 2013; Wolstenholme 1992). Complex II is encoded entirely by nuclear-encoded loci, whereas all others are mosaics of mtDNA and nuclear-encoded protein subunits. The additional mtDNA non-coding region contains: inner mitochondrial membrane and transcription factor binding sites, promoter and termination sequences, highly variable and highly conserved regions, and a 7S DNA fragment that is used to form a triple stranded mtDNA molecule (Wolstenholme 1992).

#### Accounting for shared ancestry

Principal components (PCs) are used to account for shared ancestry in the generalised linear mixed model approach used herein. To confirm that the PCs used are accounting for the variance in genotypes arising from shared ancestry, we conduct a drop test on each of the individual predictors, including individual PCs, in the full model. This test sequentially removes a single predictor from the model and tests for a reduction in model performance. For 96.5% (195 of 202) of SNPs tested, there was a statistically significant (alpha = 0.01, $p < 8.25 \times 10^{-06}$, Supplementary Table 1) drop in model performance for at least one of the PCs used. Additionally, for each of the 18 LatPrecip candidate variants, the largest reductions in model performance were observed when one of the PCs, not latitude or annual precipitation, were removed from the full model. This indicates that even for candidate variants, the majority of genotype variance is explained by shared ancestry: this is also consistent with low minor allele frequencies (typically below 0.3) for candidate variants.

#### Increase in candidate variants relative to chance

After clustering all linked SNPs (Pearson's correlation coefficient $> 0.2$) 63 distinct clusters remain. At the 1% alpha threshold we would thus expect 0.63 clusters to contain a candidate variant. We observe candidate variants within 18 of these clusters, hence representing a 28-fold enrichment in the number of candidate variants relative to the number expected by chance.

#### Stability of LatPrecip candidate variants

To ensure that signals of climate association were not artefacts of recent migrations and admixture, sequences from regions with known recent admixture (North and South America, Australia, and New Zealand) were filtered out from the dataset and the LatPrecip GLMM analysis was repeated. Of the 18 LatPrecip candidate variants reported in this manuscript, 14 were "re-identified" in the repeated analysis (Supplementary Figure 4. Of the 4 variants that were not "re-identified", the allele frequency for two variants (m.3447 and m.9950) showed a substantial decrease to near zero, leading to under-dispersion in the GLMs and the automatic exclusion of these variants from the GLM analysis following the previously described filtering stages.This is consistent with the alternative allele for these SNPs localising almost exclusively in Latin American individuals. The remaining 2 candidate variants that were not "re-identified" (m.11944 and m.14212) in the analysis without sequences from the Americas, Australia, or New Zealand remain statistically significant, however the dispersion ratio of the models for these variants increase tenfold (i.e. there is more variation than the model predicts), and therefore are also automatically filtered out of the list of candidates from the re-run analysis.

### Sequences per continent

| Continent | Number of Sequences |
|---|---|
| Europe | 8256 |
| Asia | 4491 |
| Africa | 2150 |
| Oceania | 1891 |
| South America | 1502 |
| North America | 1280 |

Table 1: Number of sequences per continent

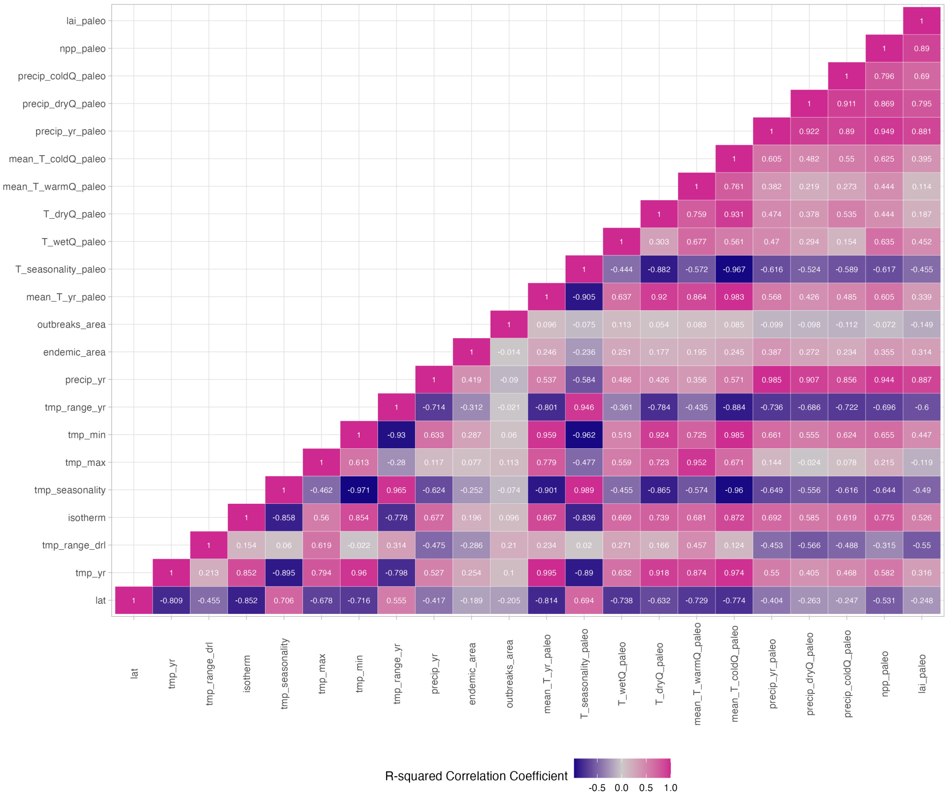


Supplementary 1 - Figure 1: **Correlations between environmental variables.** Values denote Pearson's $R^2$ correlation coefficients between variables. "_paleo" suffix denotes environmental variable is reconstructed data obtained from the pastclim R package Leonardi et al. 2023, all other environemental variables obtained from WORLDCLIM Fick and Hijmans 2017. lat: Latitude; tmp_yr / mean_T_yr: mean annual temperature (ºC); tmp_range_drl: diurnal temperature range (ºC); isotherm: isothermality; tmp_seasonality: temperature seasonality; tmp_max: maximum annual temperature (ºC); tmp_min: minimum annual temperature (ºC); tmp_range_yr: temperature range per annum (ºC); precip_yr: annual precipitation (mm); endemic_area: the number of endemic species (from WCVP) normalised by country area; outbreaks_area: the number of outbreaks reported normalised by country area; T_wetQ_paleo: mean temperature of the wettest quarter (ºC); T_dryQ_paleo: mean temperature of the driest quarter (ºC); mean_T_warmQ_paleo: mean temperature of the warmest quarter (ºC); mean_T_coldQ_paleo: mean temperature of the coldest quarter (ºC); precip_dryQ_paleo: precipitation of the driest quarter (mm); precip_coldQ_paleo: precipitation of the coldest quarter (mm); npp: Net Primary Productivity; lai_paleo: Leaf Area Index.

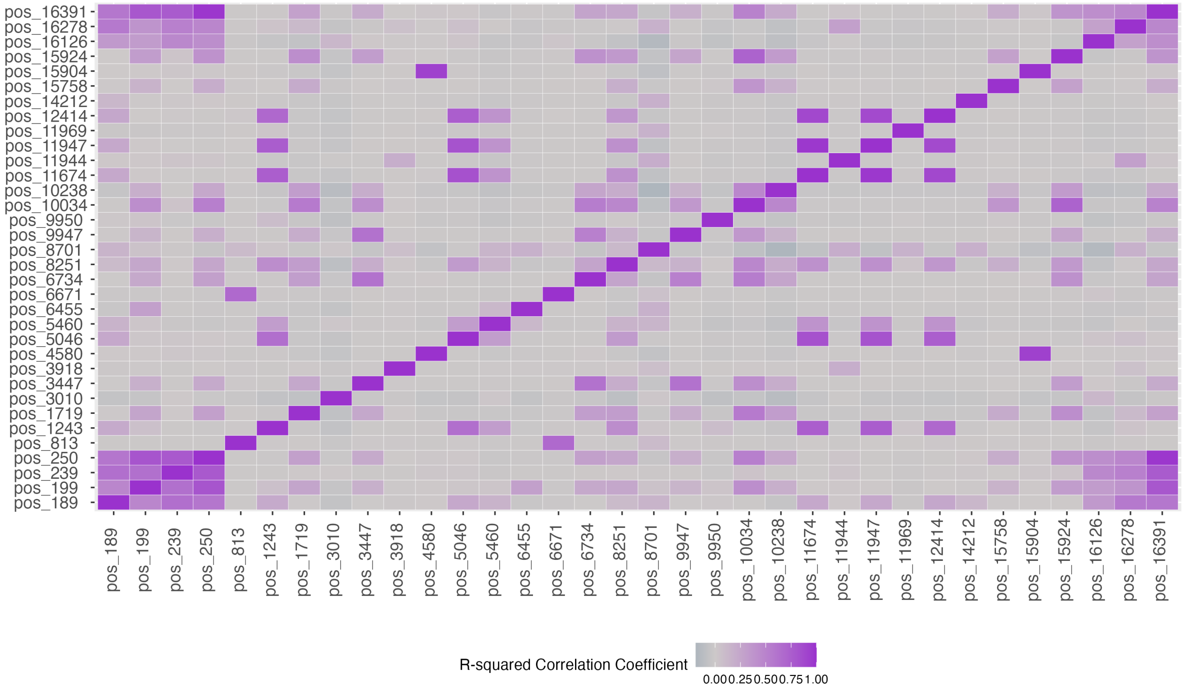


Supplementary 1 - Figure 2: **Pearson's $R^2$ correlation coefficients** between all SNPs where the LatPrecip GLMM significantly outperformed the null model. Darker purple represents a stronger correlation between SNP genotypes. SNPs with correlation coefficients above 0.2 were clustered together for downstream analysis.

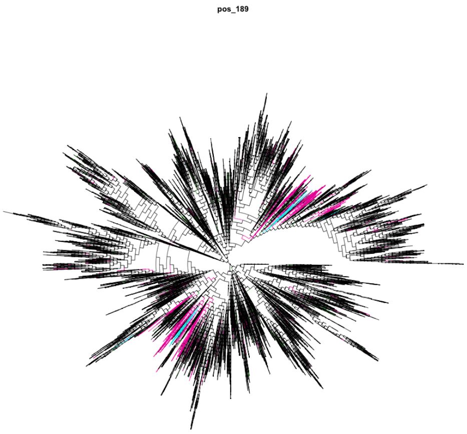


(a) m.189

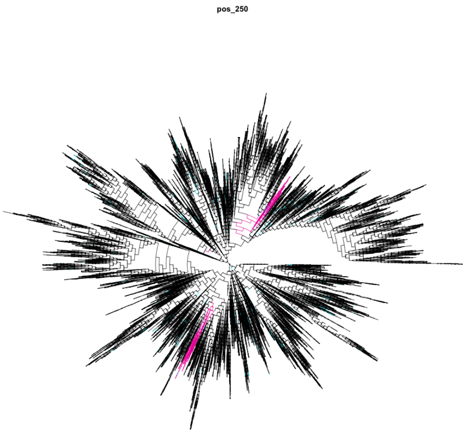


(b) .250

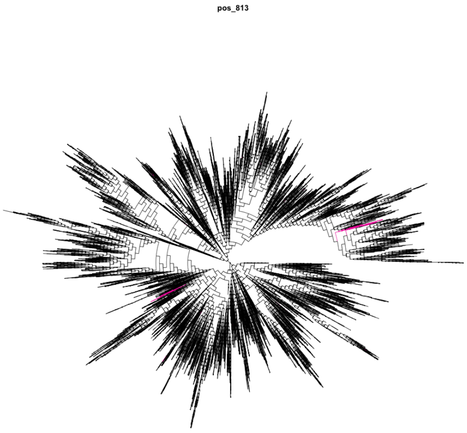


(c) m.813

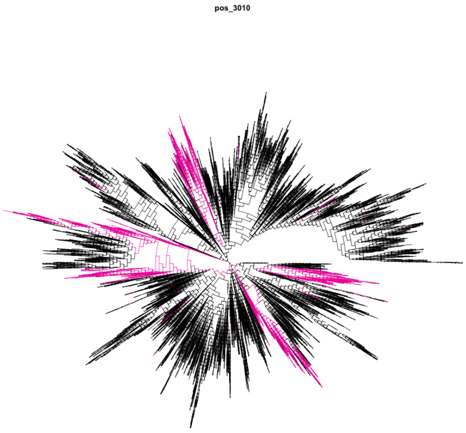


(d) m.3010

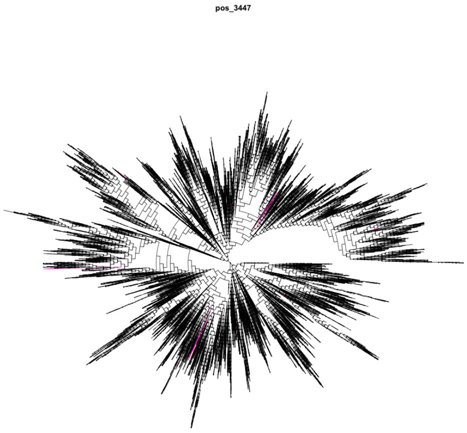


(e) m.3447

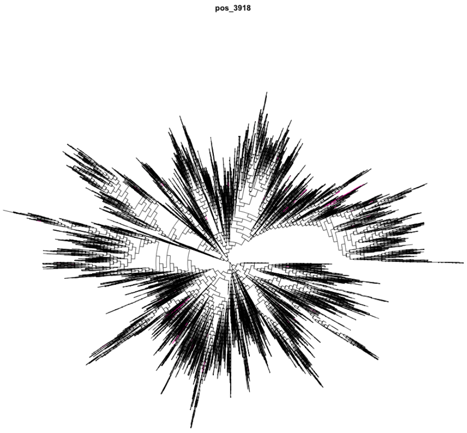


(f) m.3918

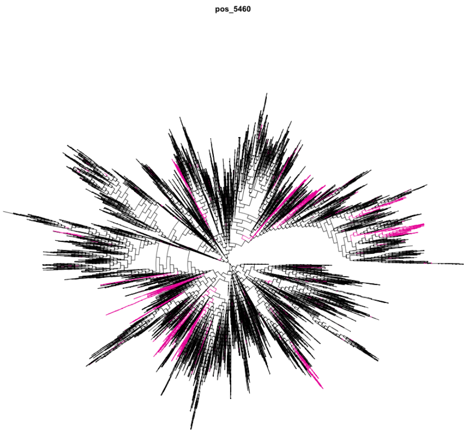


(g) m.5460

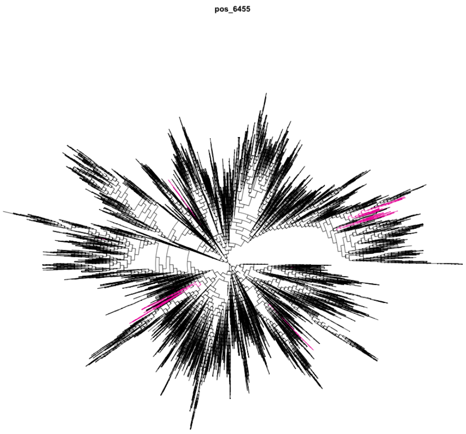


(h) m.6455

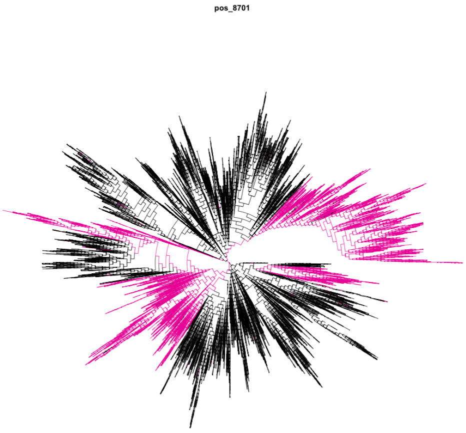


(i) m.8701

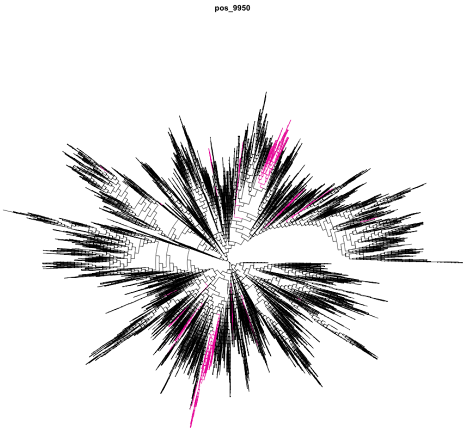


(j) m.9950

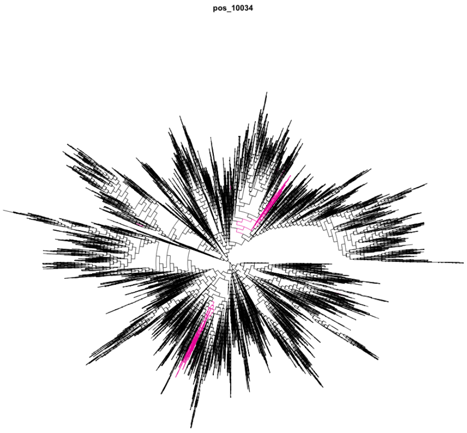


(k) m.10034

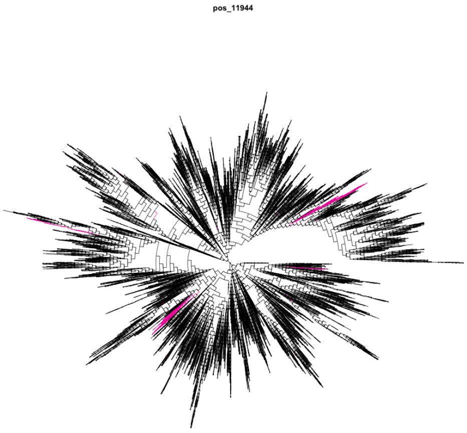


(l) m.11944

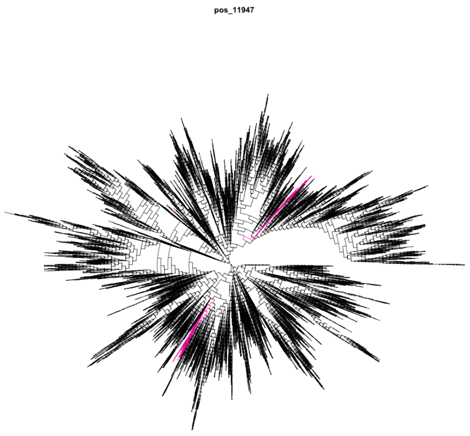


(m) m.11947

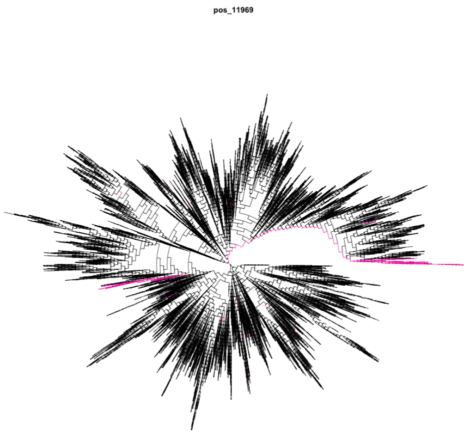


(n) m.11969

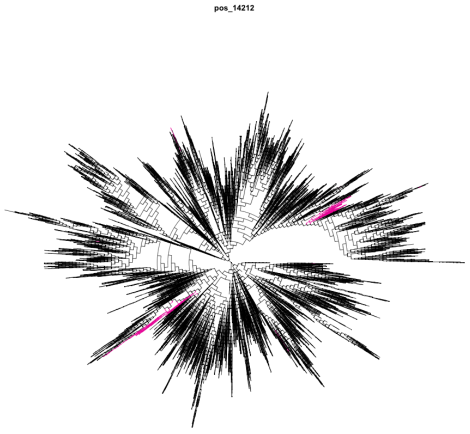


(o) m.14212

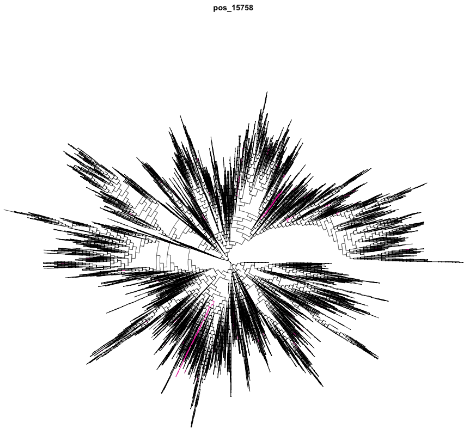


(p) m.15758

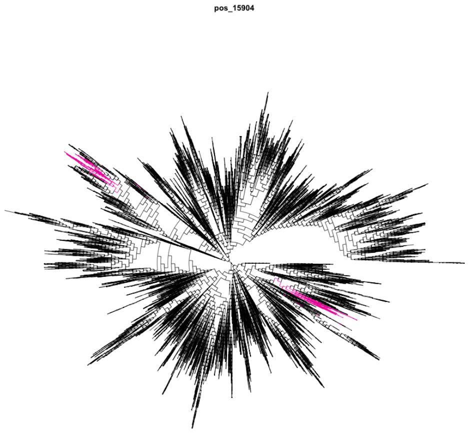


(q) m.15904

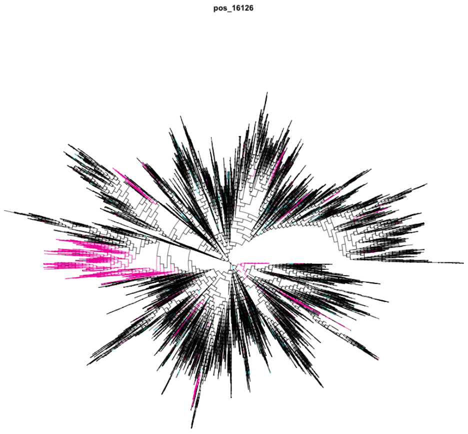


(r) m.16126

Supplementary 1 - Figure 3: **Candidate Variant Trees.** Phylogenetic trees for all 19,570 sequences used in the GLMM and TreeWAS approaches coloured (ancestral: black; derived: pink, blue) for each SNP.

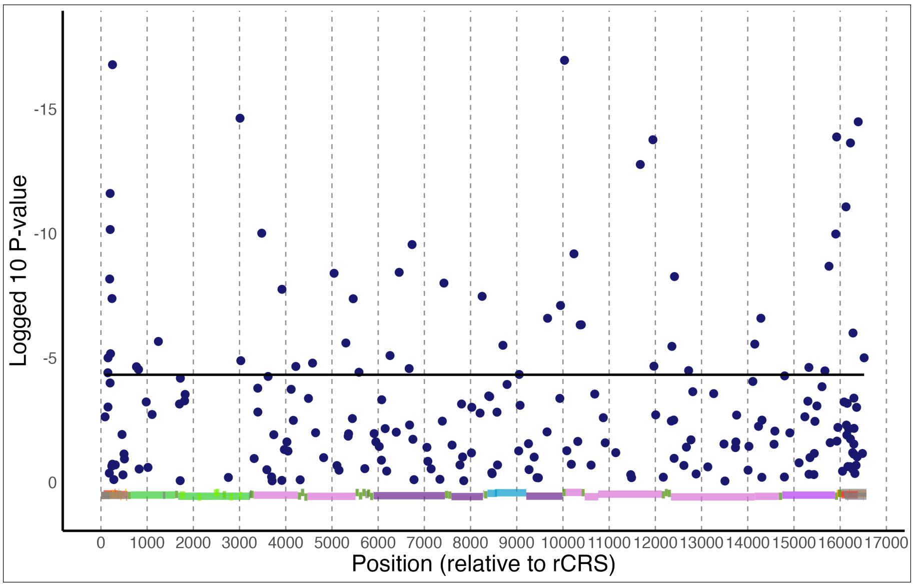


Supplementary 1 - Figure 4: **Climate associations of mtDNA variants after removing sequences from the Americas, Australia, and New Zealand.** Manhattan plot shows signals of association are not artefacts of recent migration and admixture. $\text{Log}_{10}$ p-values from likelihood ratio tests comparing GLMMs with and without climate variables (latitude and annual precipitation). Blue dots indicate a score for each of the 209 SNPs tested. The black solid horizontal line denotes the Bonferroni-corrected 1% significance threshold. Coloured blocks show loci classification (red & orange: control region; light green: rRNA subunits; dark green: tRNAs; light pink: complex I subunits; light purple: *cytB*; dark purple: complex IV subunits; blue: complex V subunits).

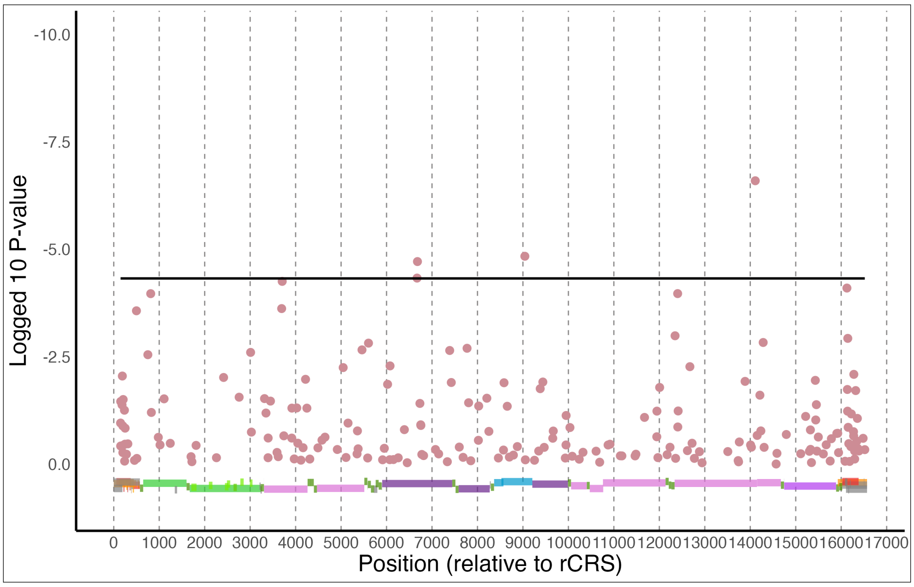


Supplementary 1 - Figure 5: **Capture of environmental variance.** $\text{Log}_{10}$ p-values from likelihood ratio tests comparing GLMMs with and without environmental PC1 relative to GLMMs with latitude and annual precipitation alongside shared ancestry. All but 4 SNPs (2 of which are correlated) fail to reach significance, indicating that latitude and annual precipitation accurately capture the environmental variance for the vast majority of SNPs, including all "LatPrecip candidate variants". Pink dots indicate a score for each of the 209 SNPs tested. The black solid horizontal line denotes the Bonferroni-corrected 1% significance threshold. Coloured blocks show loci classification (red & orange: control region; light green: rRNA subunits; dark green: tRNAs; light pink: complex I subunits; light purple: *cytB*; dark purple: complex IV subunits; blue: complex V subunits).

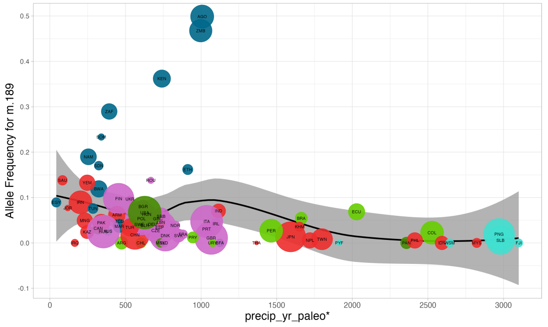
Allele Frequency for m.189
precip_yr_paleo*

(a)

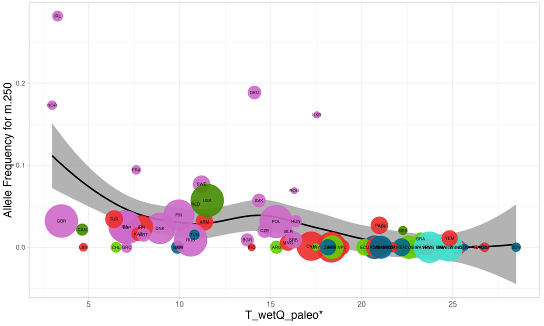
Allele Frequency for m.250
T_wetQ_paleo*

(b)

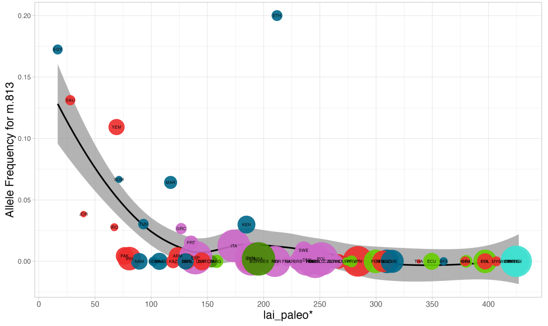
Allele Frequency for m.813
lai_paleo*

(c)

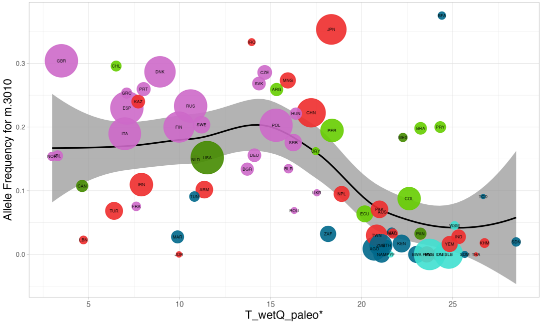
Allele Frequency for m.3010
T_wetQ_paleo*

(d)

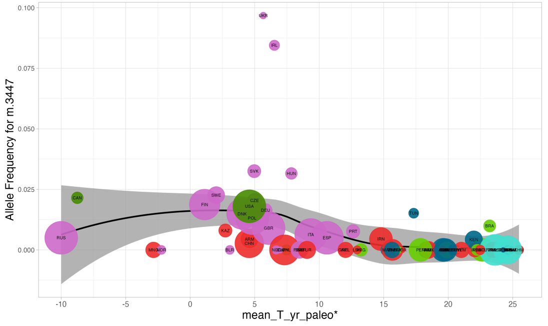
Allele Frequency for m.3447
mean_T_yr_paleo*

(e)

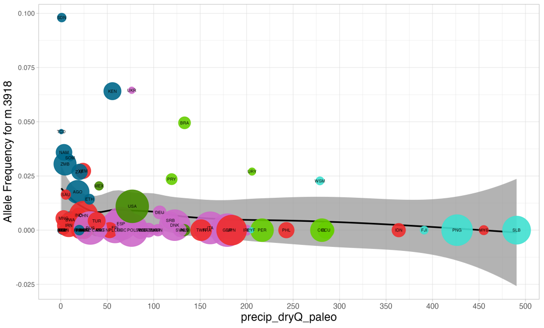
Allele Frequency for m.3918
precip_dryQ_paleo

(f)

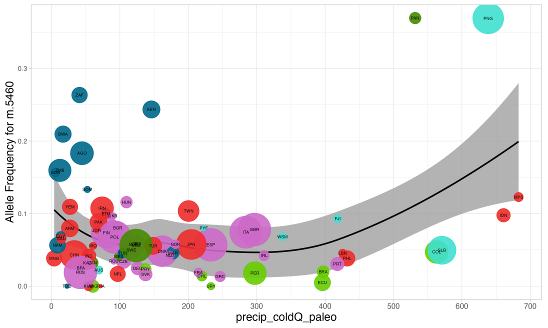
Allele Frequency for m.5460
precip_coldQ_paleo

(g)

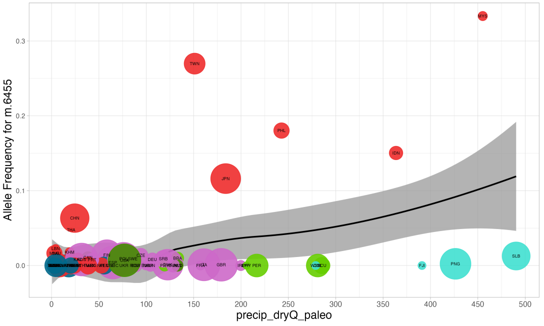
Allele Frequency for m.6455
precip_dryQ_paleo

(h)

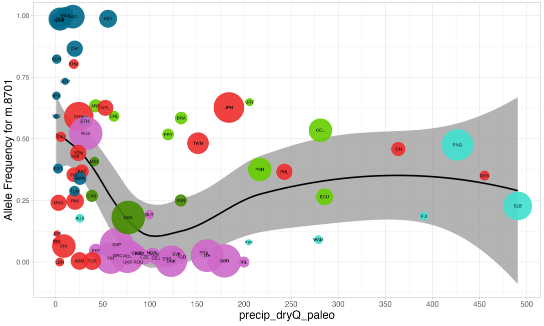
Allele Frequency for m.8701
precip_dryQ_paleo

(i)

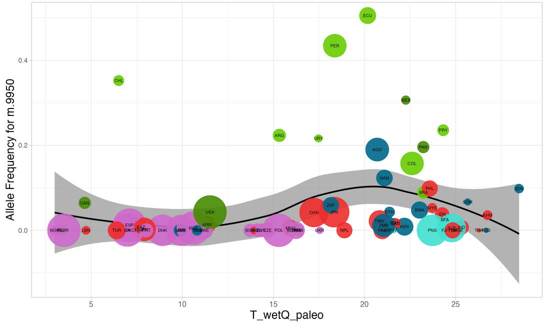
Allele Frequency for m.9950
T_wetQ_paleo

(j)

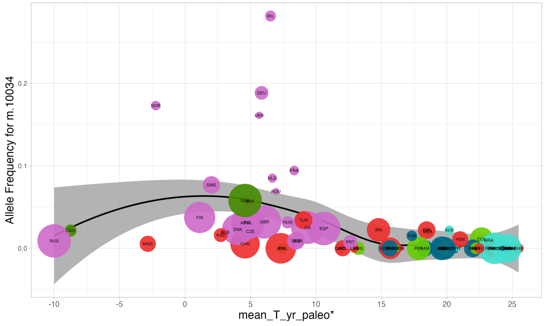


(k)

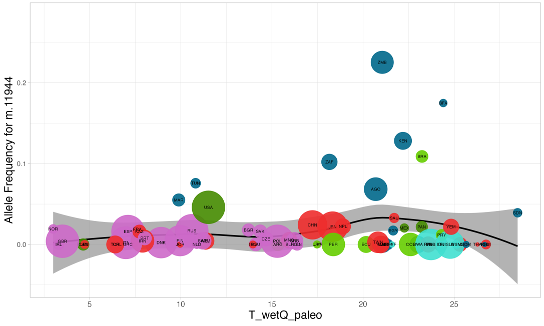


(l)

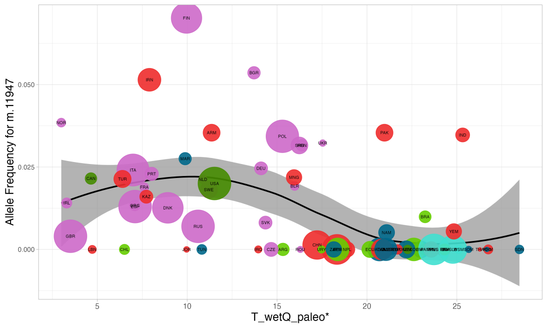


(m)

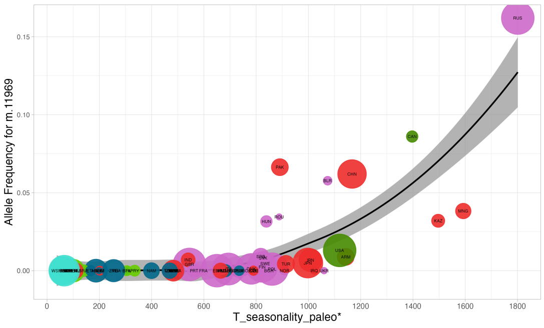


(n)

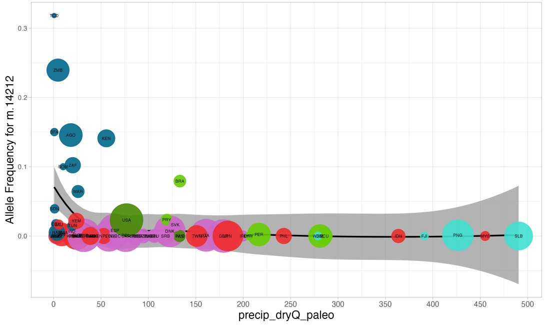


(o)

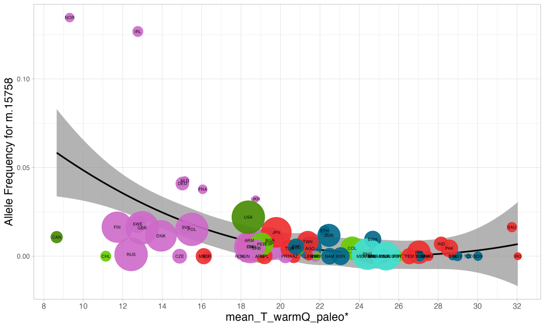


(p)

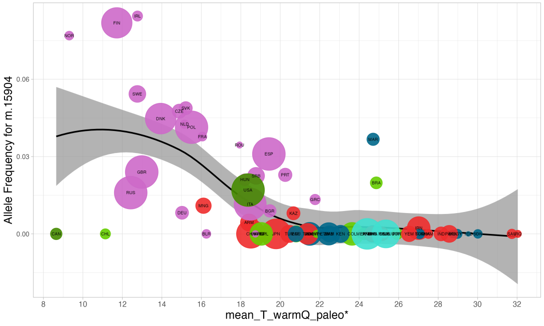


(q)

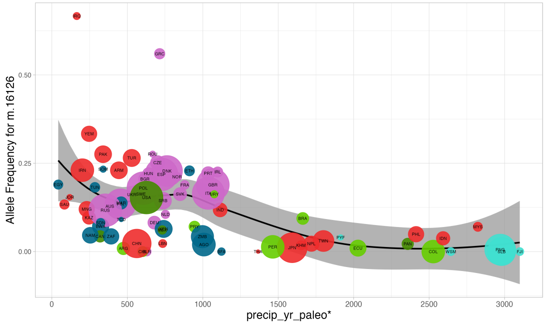


(r)

Supplementary 1 - Figure 6: **Candidate Variant Allele Frequencies** relative to paleo-bioclimatic variables. Best performing PaleoBio predictor represented. "*" denotes that the PaleoBio-GLMM significantly outperformed the null after Bonferrroni correction. Iso3 codes listed per circle. Colour represents continent: Africa (navy), Asia (red), Europe (pink), North America (dark green), South America (light green), and Oceania (turquiose). precip_yr_paleo: mean annual precipitation; T_wetQ_paleo: mean temperature of the wettest quarter (ºC); lai_paleo: Leaf Area Index; mean_T_yr_paleo: mean annual temperature (ºC); precip_dryQ_paleo: precipitation of the driest quarter (mm), precip_coldQ_paleo: precipitation of the coldest quarter (mm); precip_dryQ_paleo: precipitation of the driest quarter (mm); T_seasonality_paleo: temperature seasonality (ºC); mean_T_warmQ_paleo: mean temperature of the warmest quarter (ºC).

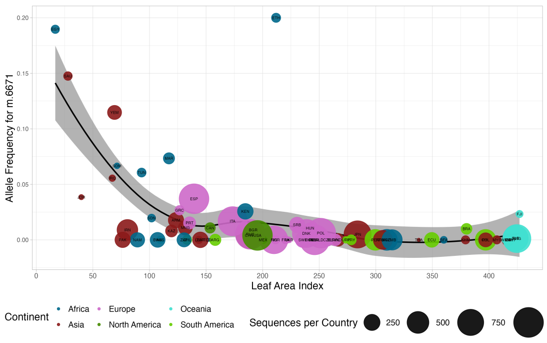


(a)

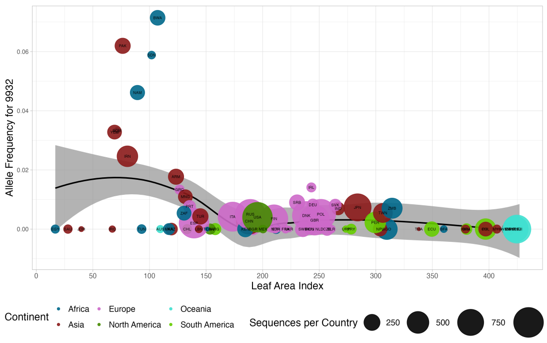


(b)

Supplementary 1 - Figure 7: **'New' PaleoGLMM Candidate Variant Allele Frequencies** relative to Leaf Area Index. m.6671 and m.9932 shown as these were not identified as LatPrecip candidate variant (see 6 for the equivalent figure for those variants). Iso3 codes listed per circle. Colour represents continent: Africa (navy), Asia (red), Europe (pink), North America (dark green), South America (light green), and Oceania (turquiose). lai_paleo: Leaf Area Index

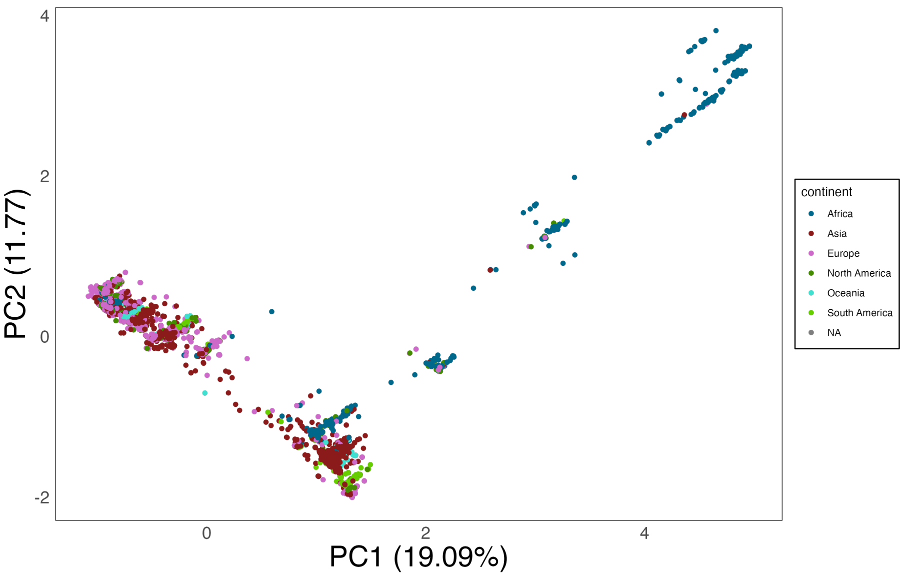


(a)

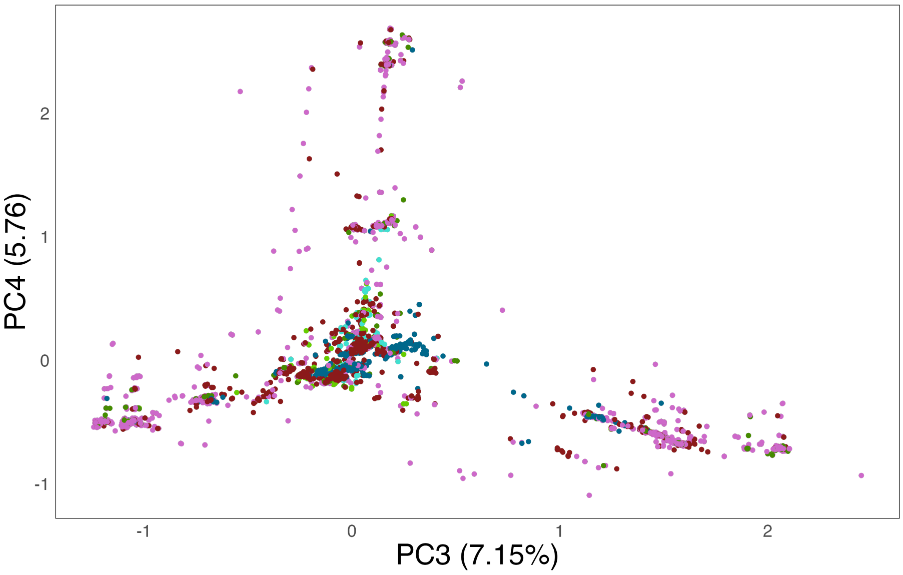


(b)

Supplementary 1 - Figure 8: **Principal Component (PC) Analysis.** PC1 vs PC2 (8a) and PC3 vs. PC4 (8b). Percentage of variance explained written in brackets. Colour denotes continent of sequence origin; Africa (navy), Asia (red), Europe (pink), North America (dark green), South America (light green), Oceania (turquoise).

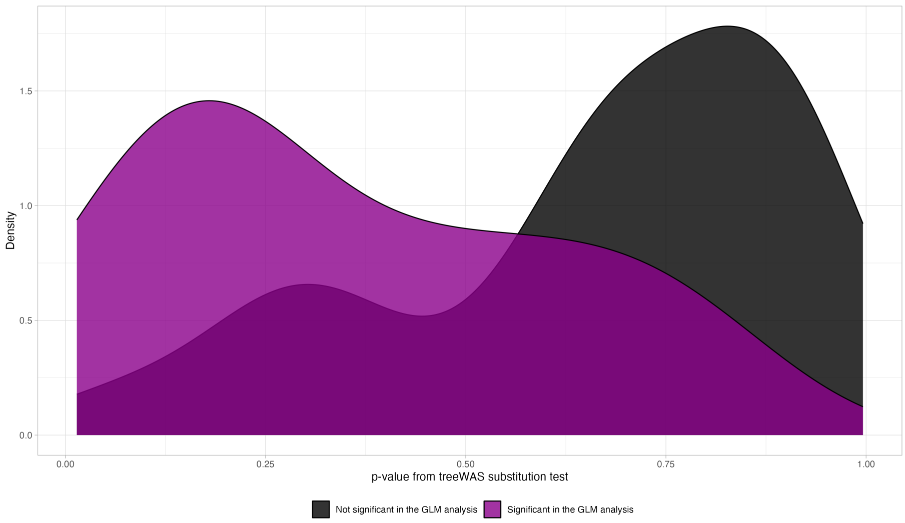


Supplementary 1 - Figure 9: **TreeWAS vs GLMM comparison.** Density plot showing the distribution of TreeWAS p-values for LatPrecip candidate variants (purple) and linked sites versus SNPs that did not correlate with LatPrecip candidate variants (black). SNPs in linkage with LatPrecip candidate variants skew towards 0.
