## Supplementary 2 for "Climate associated natural selection in the human mitochondrial genome"

### Supplementary File 2: Climate associated natural selection in the human mitochondrial genome.

Finley Grover-Thomas[1], Lucy van Dorp[1], Francois Balloux[1] , Aida M. Andrés[1], M. Florencia Camus[1]*

[1] Department of Genetics, Evolution and Environment, University College London, London, UK

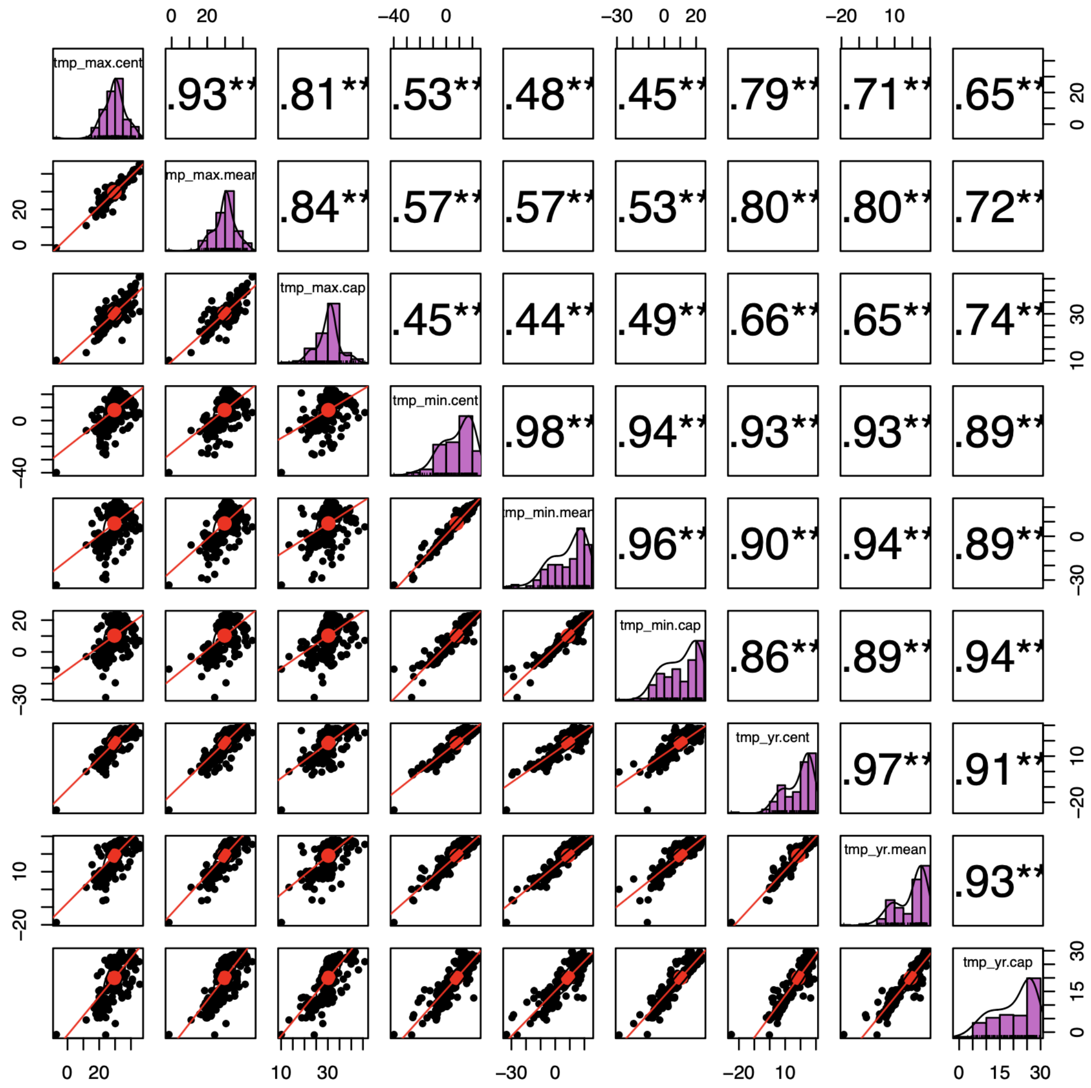


Supplementary 2 - Figure 1: **Comparison of environmental variables from mean (".mean") values vs. centroid (".cent") or capital coordinates (".cap").** Prefix denotes environmental variable. Diagonal represents histogram of a given variables' distribution. Upper triangle indicates correlation coefficients (stars denote significance), lower triangle shows scatter plots between the variables.

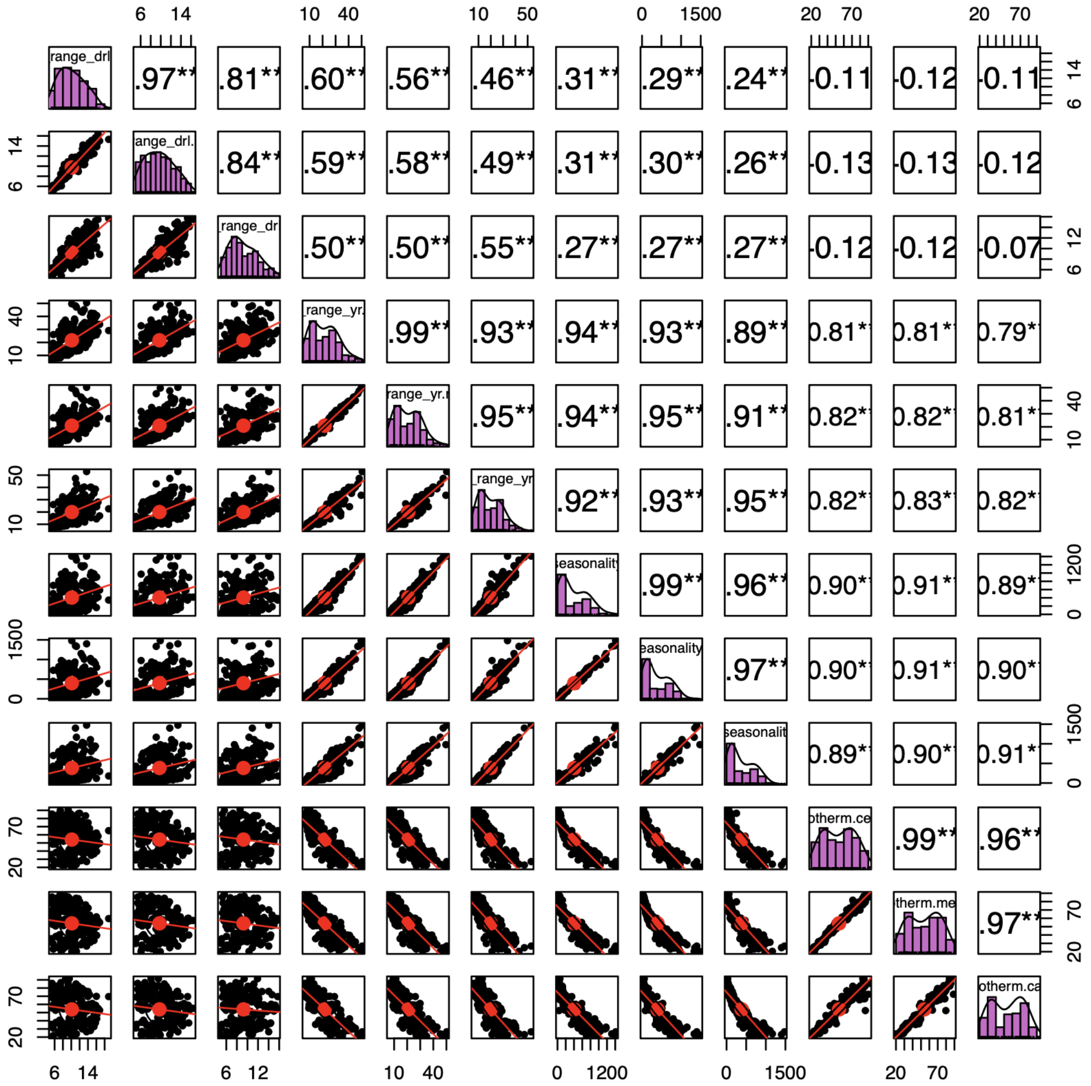

Supplementary 2 - Figure 2: **Comparison of environmental variables from mean (".mean") values vs. centroid (".cent") or capital coordinates (".cap").** Prefix denotes environmental variable. Diagonal represents histogram of a given variables' distribution. Upper triangle indicates correlation coefficients (stars denote significance), lower triangle shows scatter plots between the variables.

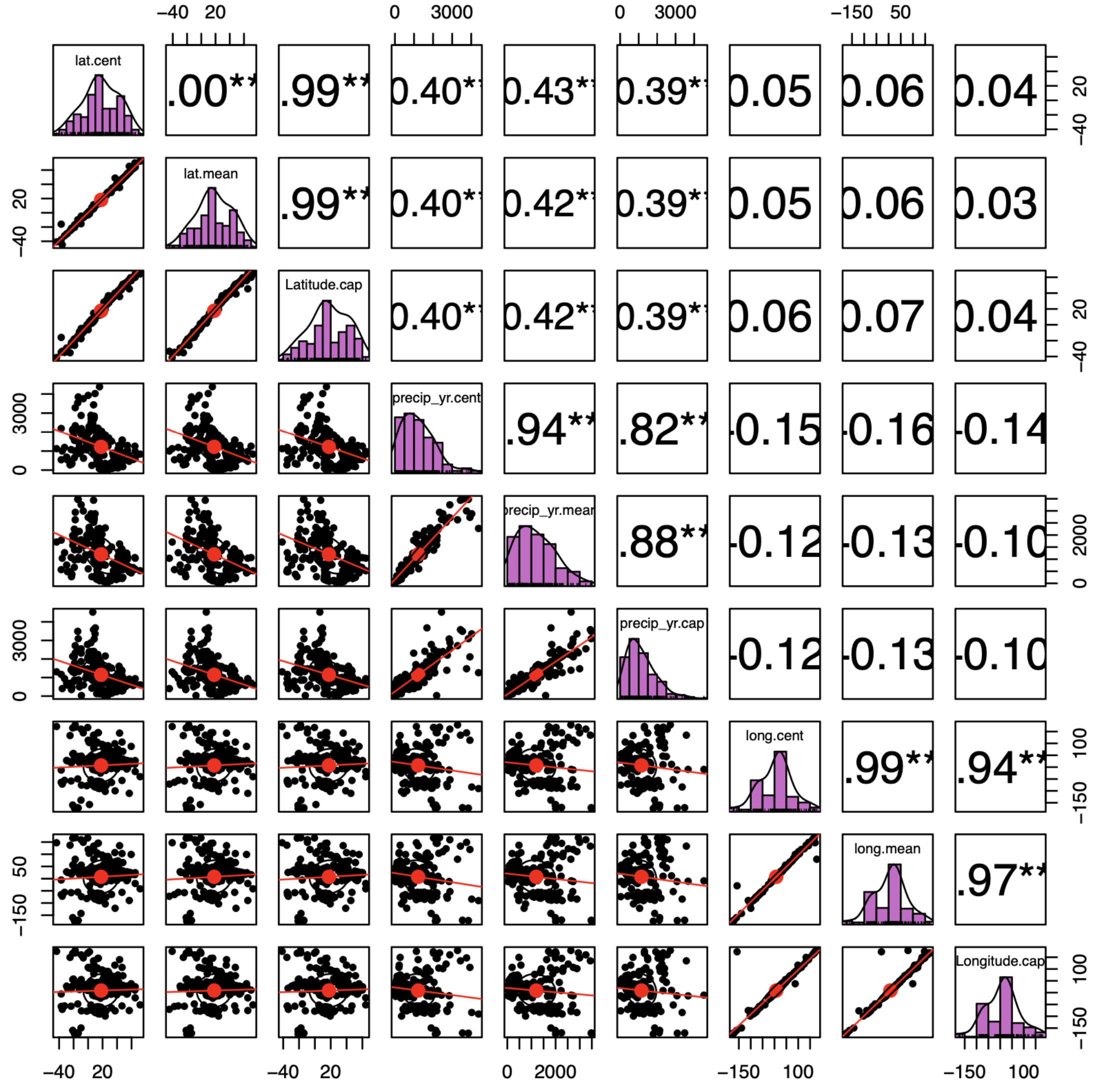


Supplementary 2 - Figure 3: **Comparison of environmental variables from mean (".mean") values vs. centroid (".cent") or capital coordinates (".cap").** Prefix denotes environmental variable. Diagonal represents histogram of a given variables' distribution. Upper triangle indicates correlation coefficients (stars denote significance), lower triangle shows scatter plots between the variables.
